# Temporal dynamics and genomic programming of plasma cell fates

**DOI:** 10.1101/2023.08.27.555004

**Authors:** Godhev Kumar Manakkat Vijay, Ming Zhou, Kairavee Thakkar, Abigail Rothrauff, Amanpreet Singh Chawla, Dianyu Chen, Louis Chi-Wai Lau, Peter Habib, Kashish Chetal, Prabal Chhibbar, Jingyu Fan, Jishnu Das, Alok Joglekar, Lisa Borghesi, Nathan Salomonis, Heping Xu, Harinder Singh

## Abstract

Affinity-matured plasma cells (PCs) of varying lifespans are generated through a germinal center (GC) response. The developmental dynamics and genomic programs of antigen-specific PC precursors remain to be elucidated. Using a model antigen, we demonstrate biphasic generation of PC precursors, with those generating long-lived bone marrow PCs preferentially produced in the late phase of GC response. Clonal tracing using scRNA-seq+BCR-seq in spleen and bone marrow compartments, coupled with adoptive transfer experiments, reveal a novel PC transition state that gives rise to functionally competent PC precursors. The latter undergo clonal expansion, dependent on inducible expression of TIGIT. We propose a model for the proliferation and programming of precursors of long-lived PCs, based on extended antigen encounters followed by reduced antigen availability.

## Introduction

Plasma cells (PCs) are considered to represent a terminally differentiated state generated by the encounter of B cells with antigens in the context of pathogens or vaccines. PCs constitutively secrete antibodies which can serve as a source of protective antibody responses^1–5^. Following antigen exposure, in the context of a T cell-dependent response, antigen-stimulated B cells interact with T follicular helper (Tfh) cells in secondary lymphoid organs (SLOs), e.g., spleen or lymph nodes, undergo clonal expansion, somatic hypermutation and affinity maturation in germinal centers (GCs), generating both memory B cells and PC precursors, the latter migrate through the bloodstream and home to the bone marrow (BM), where they undergo further maturation to terminally differentiate into PCs^6,7^. The nature of signaling pathways and transcriptional programs that result in the generation of PC precursors in SLOs that are functionally competent to migrate through the bloodstream to the bone marrow still need to be thoroughly understood.

Antiviral antibody responses can be remarkably stable in humans, lasting decades in the case of varicella–zoster and measles viruses, but are less durable for influenza viruses^8^. The durability of antibody responses to viral infections and vaccines reflects the longevity of PCs within the bone marrow (BM)^9,10^. The cellular and molecular mechanisms underlying the generation of short-lived versus long-lived PCs (SLPCs, LLPCs) are of heightened interest given the recent COVID-19 pandemic^11^. The longevity of PCs in the bone marrow could be dictated by the transcriptional programming of PC precursors emanating from the GC and/or by niches in the bone marrow that the PCs reside within.

The temporal dynamics of PC precursor generation have been inferred based on the emergence of antigen-specific PCs in the spleen as well as in the bone marrow, in the context of NP-specific B cell responses in murine models. A key study tracked the responses of NP-specific B cells in the spleen and bone marrow between 7 to 28 days post-immunization (d.p.i.)^12^. Splenic NP-specific IgG1^+^ antibody-secreting cells (ASCs) peaked at 14 d.p.i. and then declined, thereby primarily reflecting the generation of extrafollicular plasmablasts. In contrast, ASCs in the bone marrow manifested a nearly 5-fold increase between 14 and 28 d.p.i.^13^. These results suggested that bone marrow PC (BMPC) precursors are maximally generated in a temporally delayed manner during an ongoing GC response. The temporal dynamics of PC precursor generation have also been analyzed using an alternate NP-specific model. In this model, NP-reactive B cells isolated from B1-8 mice were transferred into AM14 transgenic Vk8R mice, prior to immunization with NP-CGG^12^. Two waves of ASC generation were noted in the spleen, with peaks at 11 and 38 (d.p.i.). Notably, the latter peak coincided with the maximal emergence of ASCs in the bone marrow that included LLPCs^12^. However, the nature of the PC precursors implicated by these studies and the mechanisms underlying their generation and/or expansion at later phases of the GC response remain to be delineated.

During PC differentiation, B cells undergo extensive genomic re-programming, which results in the repression of a large set of B cell genes and the activation of PC-specific as well hematopoietic progenitor and T cell genes^14,15^. This process is regulated by various transcription factors (TFs) principally, IRF4, BLIMP1, XBP1 and ATF6b^16–20^. Loss of IRF4 in PCs affects their survival and the expression of PC genes, including those required for the elaboration of the endoplasmic reticulum (ER), thereby impairing antibody secretion. BLIMP1 primarily controls the expression of components of the unfolded protein response (UPR) including the genes encoding the direct regulators of the UPR, namely the transcription factors XBP1 and ATF6b. Deletion of IRF4 also results in an increase of mitochondrial mass and oxidative phosphorylation capacity and enhanced expression of IRF8, a counteracting regulator that is expressed at high levels in GC B cells, along with BCL6, which promotes affinity maturation while antagonizing PC differentiation^21,22^. These results suggest that the induction of IRF4 could initiate the programming of PC precursors, in part via their metabolic reprogramming, as they emanate from the GC. In keeping with this possibility, NP-reactive BCL6^lo^IRF4^hi^ cells have been identified in the light zone (LZ) of the GC and suggested to represent PC precursors based on the expression of IRF4-regulated PC genes^23^. It remains to be determined if such cells are also generated at later time points in the NP-response, and if they give rise preferentially to LLPCs in the bone marrow. GC B cells receive two major signals from Tfh cells, CD40L and IL-21, that are required to generate PC precursors^23,24^. Haploinsufficiency of *Stat3* in GC B cells, a transcription factor that is activated downstream of IL-21 signaling, impairs the generation of PC precursors^23^. Furthermore, stimulation of GC B cells with CD40L and IL-21 results in an increased frequency of BCL6^lo^IRF4^hi^ cells^25^. The signaling interplay between CD40, IL-21R, and antigen recognition via the BCR, and how such stimuli dictate the generation of PC precursors as well as the fates of their PC progeny, remain to be explored. Thus, deeper analyses of the signaling and genomic regulatory mechanisms that underlie the generation of PC precursors generated during GC-dependent B cell responses will not only advance fundamental understanding of these distinctive cells but also enable their precise therapeutic manipulation so as to enhance vaccination strategies^26,27^ and target pathogenic counterparts^28–30^.

We have previously utilized scRNA-seq in conjunction with BCR-seq to track NP-specific B cell responses in GCs and the dynamic genomic states associated with class switch recombination, somatic hypermutation and affinity maturation^21,31^. We now extend this approach to the emergence of PC precursors from the GC and tracking of B cell clones in the splenic and bone marrow compartments. We demonstrate a biphasic generation of BMPC precursors, with those generating long-lived PCs preferentially produced in the late phase of GC response. Clonal tracing in spleen and bone marrow compartments, coupled with adoptive transfer experiments, reveal a novel PC transition state that gives rise to transcriptionally distinct PC precursors. The proliferation of PC precursors is dependent on the inducible expression of TIGIT. We propose a signaling model for programming of long-lived PC precursors based on extended antigen encounters followed by reduced antigen availability.

## Results

### Temporal dynamics of PC precursors generated during a GC response

To analyze antigen-specific precursors of PCs that are generated during a germinal center response in the spleen and give rise to plasma cells in the bone marrow, we designed the adoptive transfer model system schematized in Fig. 1a. Mice were immunized with NP-KLH, LPS and alum. CD138^+^ cells isolated from splenocytes (Extended Data Fig. 1a) on 21, 28, 35 and 42 d.p.i. were adoptively transferred into *Ighm*-deficient (*Ighm^tm1Cgn^*) µMT mice that lack mature B cells and PCs^32^ (Fig. 1a). Analysis of NP-specific antibody titers by ELISAs, and PCs by ELISPOTs, enabled dynamic monitoring of PC precursor activity as well as the durability of the PCs generated from them. By initiating our analysis of PC precursors at 21 d.p.i., a week after the peak of the GC response^33^, we minimized the transfer of extrafollicular plasmablasts into the recipient animals. We used CD138^+^ splenocytes for adoptive transfer as they contain both B220^+^ and B220^−^ cells and therefore represent a continuum of PC differentiation states, including the PC precursors. Notably, the CD138^+^ cells were transferred into naïve animals, thereby selecting for GC-derived PC precursors that are competent to migrate to the bone marrow and give rise to PCs in the absence of antigen.

**Fig. 1.**
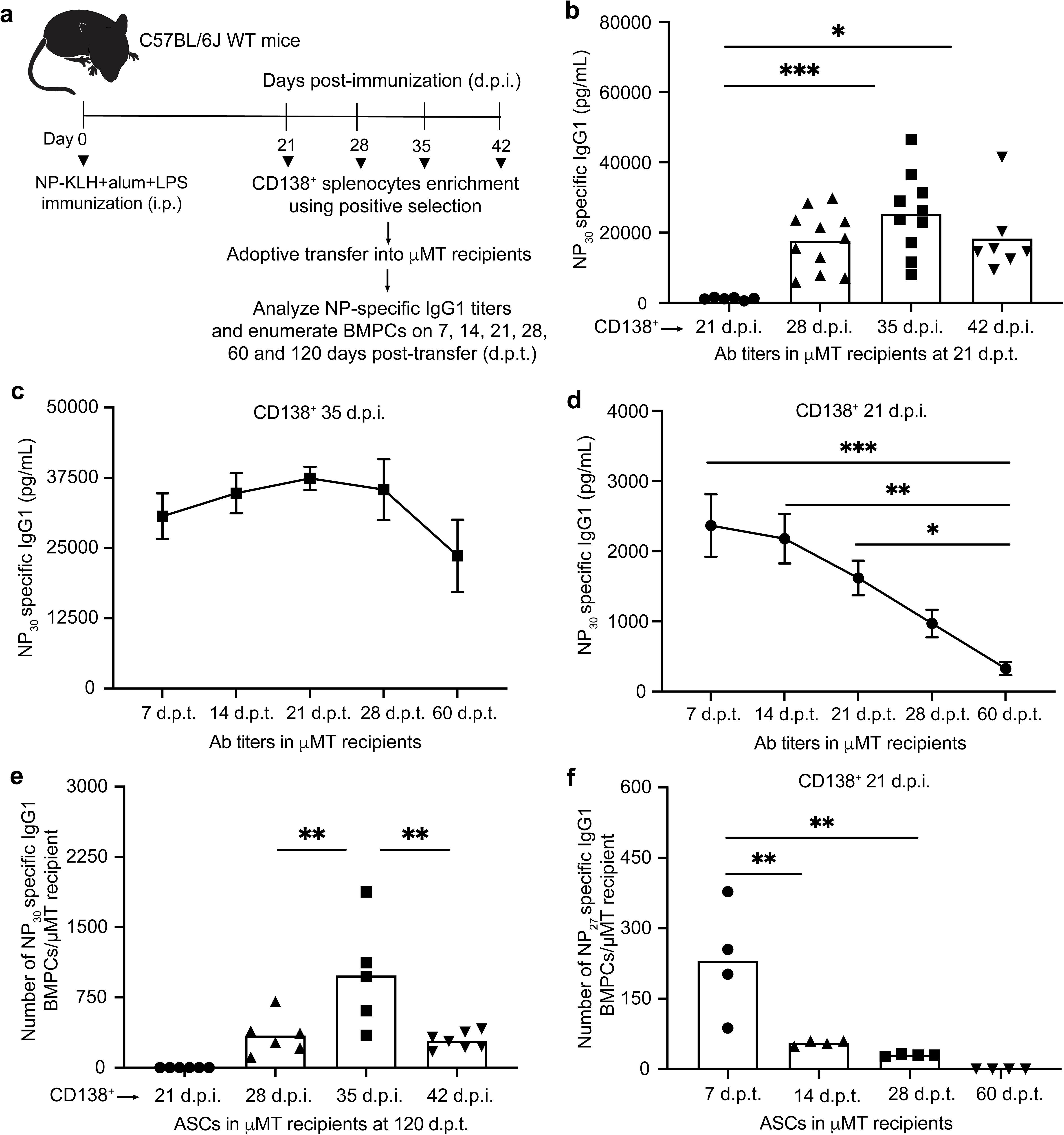
Temporal dynamics of BMPC precursors generated during a GC response. **a,** Experimental design enabling temporal analysis of BMPC precursors and the durability of their plasma cell progeny. CD138^+^ splenocytes from NP-KLH immunized mice, at indicated days post-immunization (d.p.i.), were transferred into B cell deficient µMT hosts. NP-specific antibody titers and ELISPOT analyses were performed at indicated days post-transfer (d.p.t.). **b,** Titers of NP-specific IgG1 antibodies in µMT recipients following adoptive transfer of CD138^+^ splenocytes (21, 28, 35 and 42 d.p.i.) measured at 21 d.p.t.. **c,** Titers of NP-specific IgG1 antibodies in µMT recipients (n=6) following adoptive transfer of CD138^+^ splenocytes (35 d.p.i.) at indicated d.p.t.. **d,** Titers of NP-specific IgG1 antibodies following adoptive transfer of CD138^+^ splenocytes (21 d.p.i.) in µMT recipients (n=6) at indicated d.p.t.. **e,** ELISPOT analysis of NP-specific IgG1^+^ BMPCs detected in µMT recipients following adoptive transfer of CD138^+^ splenocytes (21, 28, 35 and 42 d.p.i.) at 120 d.p.t.. **f,** ELISPOT analysis of NP-specific IgG1^+^ BMPCs detected in µMT recipients following adoptive transfer of CD138^+^ splenocytes (21 d.p.i.) at indicated d.p.t.. Each symbol represents an individual mouse (b,e,f). Data are pooled from three independent experiments shown as the mean (b,e,f) or mean ± S.E.M (c,d). Statistical significance was tested by Kruskal Wallis with Dunn’s multiple comparison test (b) or one-way ANOVA with Tukey’s multiple comparison test (c-f). *p<0.05; **p<0.01; and ***p<0.005.

Analysis of the antibody titers in the recipient animals 21 days post-transfer (d.p.t.) suggested that the peak of PC precursor activity occurred at 35 d.p.i., much later than the peak of GC response (Fig. 1b). Furthermore, whereas the antibody titers generated by CD138^+^ splenocytes isolated at 28-42 d.p.i. were durable (60 d.p.t.) those generated by CD138^+^ splenocytes isolated at 21 d.p.i. waned substantially (Fig. 1c,d, Extended data Fig. 1b,c). This suggested that PC precursors generated earlier in the GC response give rise to short-lived PCs, whereas those generated later give rise to longer-lived PCs. To directly test for the generation of PCs in the bone marrow and their longevity, we performed ELISPOT analyses in the recipient µMT mice at varying d.p.t.. As predicted by the antibody titers, the largest number of NP-specific ASCs were detected using PC precursors isolated at 35 d.p.i. (Fig. 1e). Importantly, in keeping with our expectation from the durability of antibody responses, these ASCs represented long-lived BMPCs as they were detectable at 120 d.p.t. (Fig. 1e). In contrast, such long-lived BMPCs were not detectable in adoptive transfers using PC precursors isolated at 21 d.p.i. We reasoned that splenic precursors generated at 21 d.p.i. may preferentially give rise to short-lived BMPCs. To test this possibility, we analyzed µMT recipients adoptively transferred with CD138^+^ cells from 21 d.p.i. at 7-60 d.p.t. for the durability of NP-specific PCs in the bone marrow. Notably, PCs were detectable as early as 7 d.p.t. but declined by 14 d.p.t. and were undetectable at 60 d.p.t. (Fig. 1f). These results demonstrate that SLPC precursors are generated earlier during a GC response, whereas their LLPC precursors are generated later. They suggest a temporal shift in the developmental programming of PC precursors, emanating from the GC, which results in BMPCs with longer durability at later times in the response.

### scRNA-seq reveals novel PC progenitor state and divergent PC clusters

To genomically delineate intermediates in the specification of BMPC precursors, we initially performed scRNA-seq and BCR-seq on splenic CD138^+^ splenocytes isolated at 35 d.p.i. in the GC response. This timepoint was selected as it represented maximal BMPC precursor activity based on the adoptive transfer experiments (Fig. 1). Computational analysis of the scRNA-seq dataset using the ICGS2 pipeline^34^ revealed cell clusters that could be annotated based on the expression of biologically informative marker genes (Fig. 2a, Supplementary Table 1, Methods). The appended ICGS2 viewer link facilitates user selected queries of the dynamic gene expression programs across various splenic B cell states at 35 d.p.i. (http://www.altanalyze.org/ICGS/Public/Splenic_GC_B_and_Plasma_Cell_Compartments/User.php). Within the B cell clusters, LZ and DZ B cells could be delineated from other B cells based on differential expression of a set of genes including, *Aicda*, *Bcl6* and *Basp1* (Fig. 2b, Extended Data Fig. 2a). GC B cells could be further resolved as light zone (LZ) and dark zone (DZ) subclusters based on differential expression of cell cycle regulators including *Foxm1*, *Ccnb1,* and *Plk1* (Fig. 2b, Extended Data Fig. 2a.). Plasma cell states, displaying lowered expression of B cell genes such as *Cd19*, *Pax5*, *Ptprc* and increased expression of *Irf4*, *Prdm1* and *Xbp1*, were resolved into three clusters that were demarcated by the differential expression of *Tigit*, *Slpi,* and *Lag3*, respectively (Fig. 2b and Extended Data Fig. 2b,c). ICGS2-based clustering revealed a transitional PC cell state, hereafter termed PC progenitor, as these cells retained robust expression of B-lineage specific genes but appeared to be inducing expression of *Irf4*, *Prdm1* and a large set of PC genes (Fig. 2b, Extended Data Fig. 2b). To examine developmental relationships between the PC progenitors and the cells contained in the GC as well as PC clusters, we performed single-cell trajectory analysis using Monocle2^35^. The various single-cell clusters could be organized into a continuous, well populated path with a branch at one end (Fig. 2c). GC B cells were clustered at the non-branching end of the trajectory whereas the PC clusters were positioned at the other end, primarily before and after the branchpoint. Notably, PC progenitors were dispersed along the trajectory between the GC B cells and PCs. Pseudotime analysis of the Monocle2 generated trajectory, with the non-branching end serving as the origin, aligned with the inference that PC progenitors arise from GC B cells and generate distinct types of plasma cells (Extended Data Fig. 2d). Intriguingly, the *Tigit* and *Slpi* PCs appeared to represent one terminus and the *Lag3* PCs another terminus in the trajectory (Fig. 2c). To further substantiate the developmental trajectory we performed analyses of the scRNAseq dataset using scVelo^36^. This analysis focused on the dynamics of spliced and unspliced transcripts. It reinforced the inference that PC progenitors arise from GC B cells and give rise to the various PC states (Fig. 2d). Accordingly, the PC progenitors manifested a lower PC gene signature score compared with their PC counterparts (Fig. 2e). Intriguingly, the PC compartment also displayed a higher mitotic gene expression score than the PC progenitors, suggesting that differentiated cells within the compartment undergo cell division after cell fate specification (Fig. 2f).

**Fig. 2.**
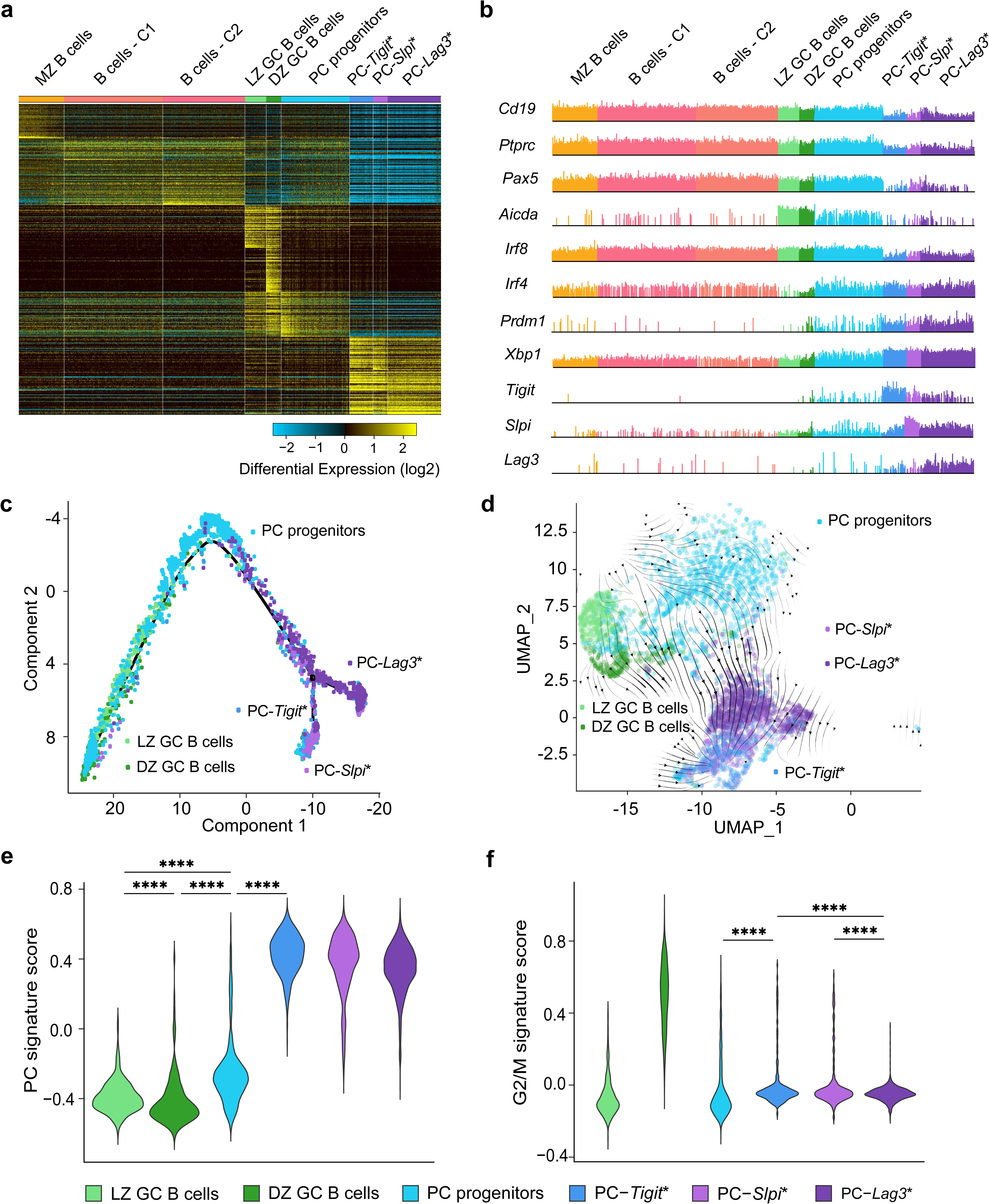
ScRNA-seq analysis of PC genomic states and trajectories. **a,** Heatmap generated using cluster-specific marker genes delineated by the MarkerFinder algorithm in AltAnalyze **(Methods)** for CD138^+^ splenocytes isolated at 35 d.p.i. (Fig. 1a) and profiled using 5’-end droplet based scRNA-seq. Columns in heatmap represent cells (n=8,813); rows represent markerfinder genes (n=413). Cell clusters were generated using ICGS2 in AltAnalyze **(Methods)** with inferred genomic states annotated on the top. MZ B cells, marginal zone B cells; B cells - C1, B cells – cluster 1; B cells - C2, B cells – cluster 2; LZ GC B cells, light zone germinal center B cells; DZ GC B cells, dark zone germinal center B cells; PC progenitors, plasma cell progenitors; PC-*Tigit*\*, PCs expressing highest *Tigit*; PC-*Slpi*\*, PCs expressing highest *Slpi*; PC-*Lag3*\*, PCs expressing highest *Lag3*. **b,** Comb plots displaying the incidence and amplitude of indicated genes in each cluster as in **a**. **c,** Developmental trajectory of GC B cells and plasma cells constructed by Monocle 2 **(Methods)**. **d,** Developmental trajectory of GC B cells and plasma cells constructed by scVelo on 2-D UMAP space **(Methods)**. **e,** Violin plots displaying the PC gene signature score **(Methods)** for indicated cell clusters. **f,** Violin plots displaying the G2/M signature score **(Methods)** for indicated cell clusters. Each dot (c,d) represents an individual cell. Statistical significance was tested by Benjamini-Hochberg for multiple test corrections (e,f). **p<0.01; ***p<0.005 and ****p<0.0001.

### Tracing antigen-specific PC states emanating from the germinal center

Given the dynamic expression of the mitotic gene module in the PC progenitors and their differentiating progeny (Fig. 2f), we sought to track PC clones using BCR-seq. To enable the analysis of antigen-specific cells responding to NP-KLH, we first tabulated *IGHV* sequences that dominate the NP-specific GC B cell response at 14 d.p.i.^37^ (Extended Data Fig. 3a,b). The top 25 NP-specific *IGHV* genes, based on their frequencies of occurrence in the NP-specific GC B cell compartment at 14 d.p.i., were used to track clones later in the GC response. Next, we delineated the top 50 *IGHV* genes that were observed at 35 d.p.i. in the CD138^+^ splenocyte compartment (Extended Data Fig. 3c and Supplementary Table 2). Finally, by generating a comparable scRNA-seq and BCR-seq dataset for CD138^+^ splenocytes at 21 d.p.i., we were able to identify NP-specific *IGHV* genes (19/25) that were prevalent and shared in both the 21 and 35 d.p.i. CD138^+^ splenocytes (Extended Data Fig. 3c,d). *IGHV* sequences shared between 21 and 35 d.p.i. cells that were not NP-specific were considered KLH-specific sequences (Extended Data Fig. 3c,d and Supplementary Table 2). Using the combined *IGHV* sequences, we annotated antigen-specific cells in the 35 d.p.i. dataset. Importantly, the selected cells exhibited the spectrum of genomic states observed with the total CD138^+^ splenocytes (Fig. 3a) and were organized continuously along the Monocle2 developmental trajectory derived using the total CD138^+^ splenocytes (Fig. 3b). To analyze and trace NP- and presumptively KLH-specific clones across genomic states, we identified cells that contained identical V(D)J and VJ gene rearrangements in their heavy and light chain immunoglobulin loci, respectively (Supplementary Table 3, Methods). Analysis of clones at 35 d.p.i. expressing the NP-specific *IGHV*1-72*01 and *IGHV*1-53*01 genes demonstrated that PC progenitors were clonally related to cells in the three PC clusters and arose from GC B cells (Fig. 3c,d and Supplementary Table 3). The latter conclusion was reinforced by the analysis of somatic hypermutations in cells harboring the dominant NP-specific *IGHV*1-72*01 (Extended Data Fig. 3e). This analysis revealed that PC progenitors and their differentiating counterparts accumulated somatic hypermutations, which increased in frequency from 21 d.p.i. to 35 d.p.i. and also evidenced affinity maturation, consistent with their GC origin (Extended Data Fig. 3f). We note that CellHarmony^38^ was used to align the single cell genomic states of the CD138^+^ splenocytes at 21 d.p.i. with their counterparts at 35 d.p.i., using the latter as the reference set (Extended Data Fig. 3g). This enabled us to reveal differentially expressed genes (DEGs) within antigen-specific PC progenitors and their differentiated progeny at 21 d.p.i. versus 35 d.p.i. (Extended Data Fig. 3h). Notably, genes involved in DNA replication were increased in their expression in PC progenitors wheras those involved in focal adhesion and ribosomal biogenesis were upregulated in the *Tigit* PCs in comparing equivalent cell states at 35 d.p.i. versus 21 d.p.i. (Fig. 3e,f, Supplementary Table 4). The two sets of DEGs suggested quantitative (cell numbers) as well as qualitative changes (cell states) in the PC precursor compartment that might account for the increase in PC precusor activity at 35 d.p.i. as well as the preferential generation of LLPCs at the later timepoint in the NP-KLH GC response (Fig. 1b,e). Given the demonstration of expanded antigen-specific clones in the splenic PC progenitors and their progeny (Fig. 3c,d) we examined their size distributions as a function of their genomic states (Fig. 3g). The analysis revealed increased sizes of clones in the PC clusters compared with the progenitors. This observation was in keeping with the higher mitotic gene signature scores of cells in the PC clusters, in relation to their progenitors (Fig. 2f) and suggested that differentiating PCs undergo proliferative expansion after their fate specification from PC progenitors that emanate from the germinal center.

**Fig. 3.**
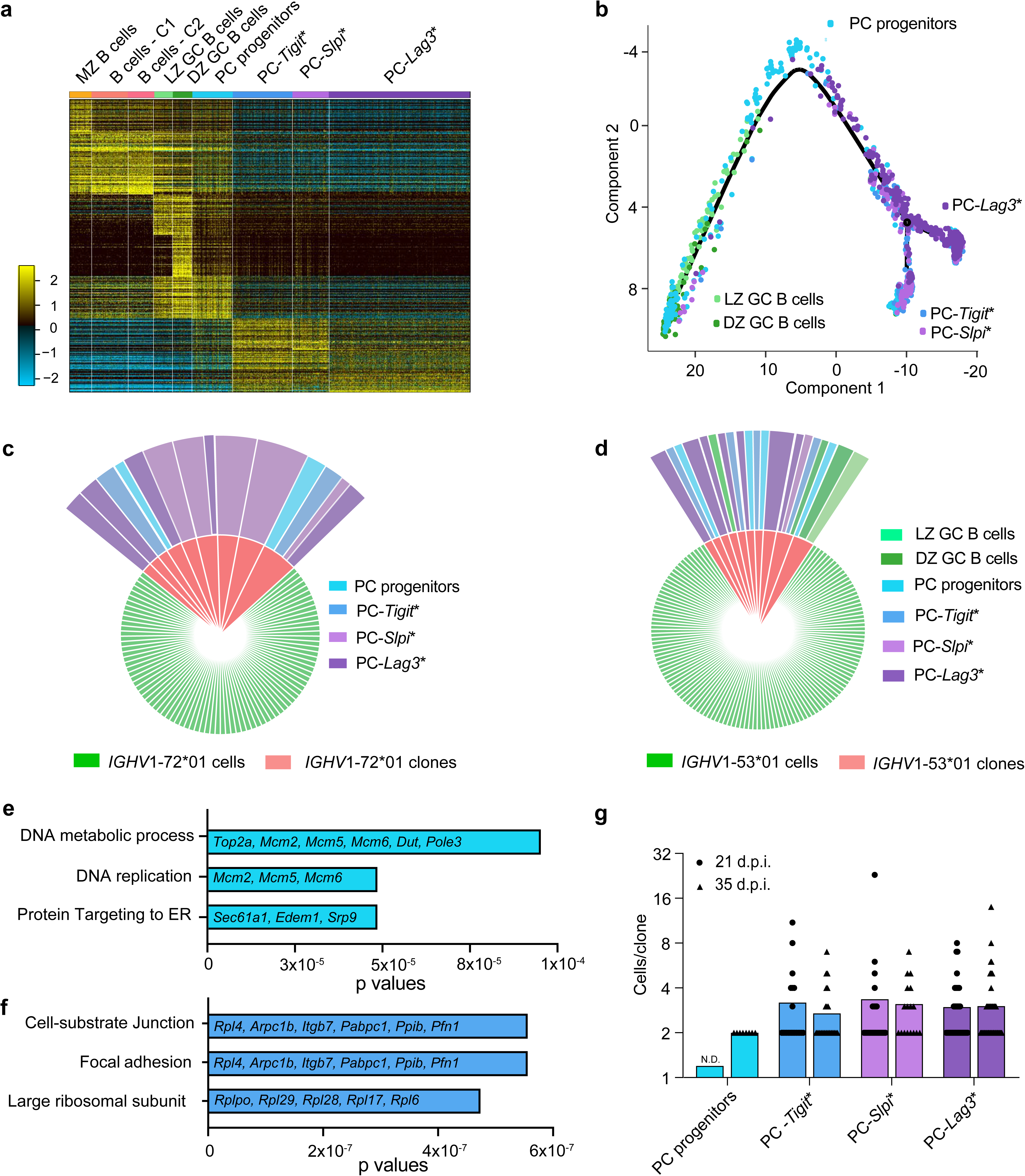
Delineation of antigen-specific PC genomic states and clonal tracking. **a,** Heatmap derived from Fig. 2a by filtering antigen-specific cells based on BCR-seq **(Methods)**. Columns and rows in heatmap represent cells (n=2,613) and MarkerFinder genes (n=413), respectively. **b,** Developmental trajectory analysis of the antigen-specific cells based on Monocle 2. Cells are color coded based on ICGS2 cluster identity as in Figure 2a. **c,d,** Pie-charts displaying the NP-dominant *IGHV*1-72*01 (V_h_186.2) **(c)** and *IGHV*1-53*01 clones **(d)**, colored by the ICGS2 cluster identities manifested within each clone. Green sectors are cells that express the rearranged *IGHV*1-72*01 or *IGHV*1-53*01 gene whereas red sectors are *IGHV*1-72*01 or *IGHV*1-53*01 clones that bear identical V(D)J rearrangements in their heavy and light chain loci. Outermost sectors display the ICGS2 cluster identities of cells within the clones. **e,** Bar plot displaying the key gene ontology pathways associated with the upregulated genes in PC progenitors (35 vs. 21 d.p.i.). **f,** Bar plot displaying the gene ontology pathways associated with the upregulated genes in PC-*Tigit*\* cluster (35 vs. 21 d.p.i.). **g,** Bar plot displaying the number of cells per clone in the indicated genomic states at 35 or 21 d.p.i.. Each dot (b) represents an individual cell. Selected genes within gene ontology pathways are displayed (e,f). Each symbol (g) represents a clone within the indicated genomic state. N.D. not detectable (g).

### Functional and genomic analyses of GC-derived PC precursors

The coupled scRNA-seq and BCR-seq analyses of antigen-specific PC cell states and their trajectories, during a GC response, revealed a novel progenitor state as well as significant clonal expansion of differentiated PC progeny. However, the nature of the BMPC precursors and their genomic state(s) within the developmental trajectory, remained to be delineated. We sought to address this question using two complementary experimental approaches (i) adoptive transfer of FACS-purified splenic CD138^+^ subsets that were genomically distinguishable and (ii) clonal tracing of antigen-specific CD138^+^ cells within splenic and bone marrow compartments of NP-KLH immunized mice. Based on prior analysis of PC precursors^13^ in the bone marrow, we utilized B220, CD138, CD44, and CD11a to resolve the splenic CD138^+^ cellular compartment (Extended data Fig. 4a). Given the dynamic expression of B220 in the PC differentiation trajectory, we purified B220^+^CD138^+^CD44^+^CD11a^+^, B220^int^CD138^+^CD44^+^CD11a^+^ and B220^−^CD138^+^CD44^+^CD11a^+^ splenic subsets from immunized mice at 35 d.p.i. and analyzed their genomic states by scRNA-seq (Fig. 4a,b and Extended data Fig. 4b). cellHarmony analysis revealed that the PC-*Lag3*\* cells were enriched within the mature B220^−^CD138^+^CD44^+^CD11a^+^ subset, whereas PC-*Tigit*\* and PC-*Slpi*\* cells predominated in the B220^int^CD138^+^CD44^+^CD11a^+^ subset. In contrast, PC progenitors were detected exclusively in an activated subset of B cells that were B220^+^CD138^+^CD44^+^CD11a^+^. ELISPOT analysis of these three subsets along with control cells suggested that the ability to secrete antibodies was acquired during the PC progenitor state, consistent with the upregulation of the PC gene module (Fig. 4c). To functionally assess which of those subsets contained BMPC precursors, we adoptively transferred the FACS-purified cells into µMT mice. NP-specific IgG1 titers, measured on 21 d.p.t., were substantially higher (>10-fold) in µMT recipients that were reconstituted with the B220^int^CD138^+^CD44^+^CD11a^+^ and B220^−^ CD138^+^CD44^+^CD11a^+^ subsets compared to the B220^+^CD138^+^CD44^+^CD11a^+^ counterparts (Fig. 4d). ELISPOT analysis of the bone marrow (60 d.p.t.) demonstrated that NP-specific IgG1^+^ ASCs were detected in comparable numbers between the B220^int^CD138^+^CD44^+^CD11a^+^ and B220^−^CD138^+^CD44^+^CD11a^+^ subsets. However, no such BMPCs were observed with the B220^+^CD138^+^CD44^+^CD11a^+^ cells (Fig. 4e). These results demonstrated that PC precursors, that are functionally competent to migrate and generate LLPCs in the bone marrow, are contained within three genomically distinct splenic PC clusters. Furthermore, the results distinguished PC progenitors from BMPC precursors as the former appeared to have acquired ASC function but not bone marrow homing/survival capabilities.

**Fig. 4.**
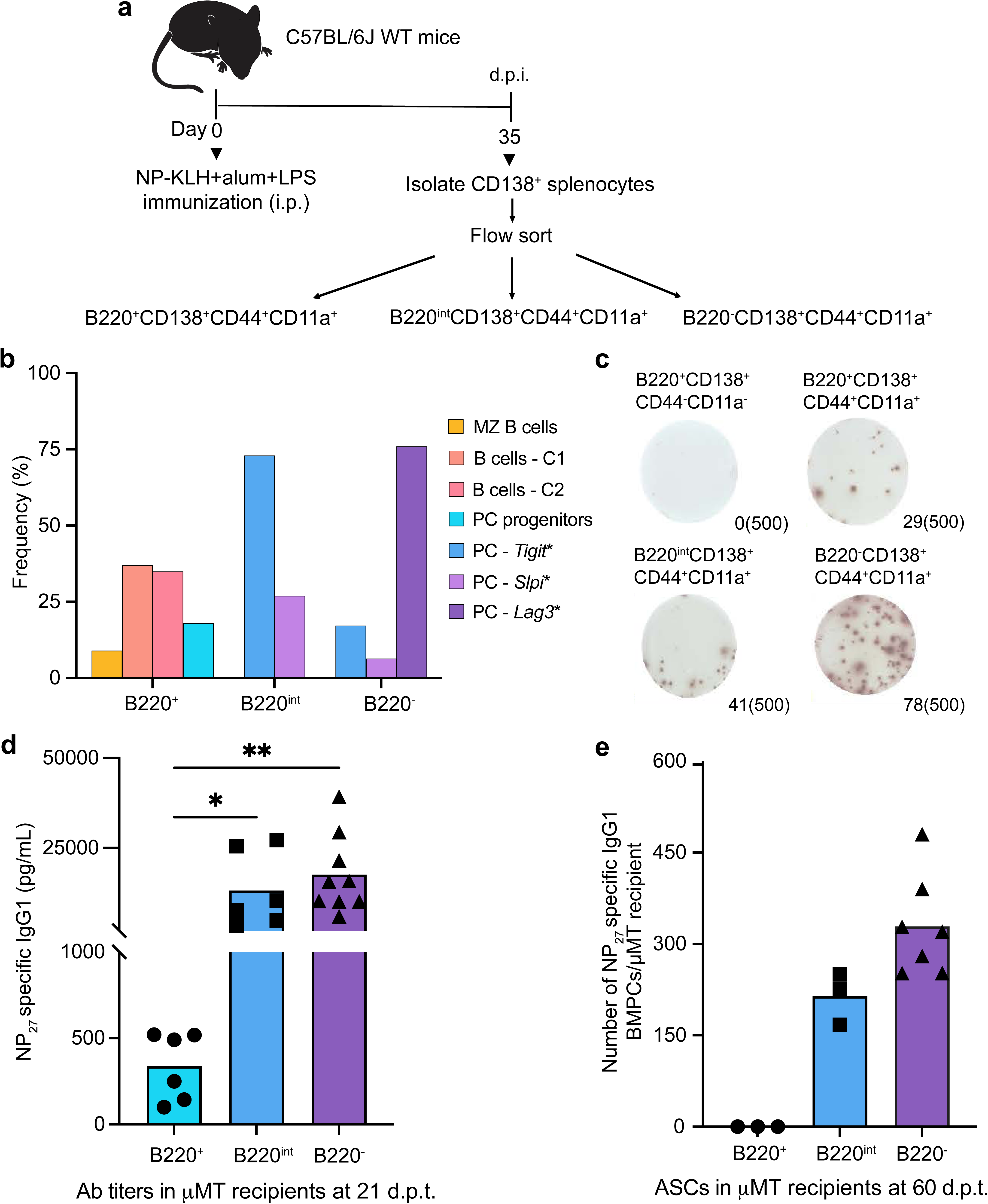
Functional and genomic analysis of BMPC precursors within splenic CD138^+^ subsets. **a,** Experimental design involving designated splenic CD138^+^ subsets isolated by flow cytometry from NP-KLH immunized mice (35 d.p.i.) and analyzed by scRNA-seq as well as by adoptive transfer into B cell deficient µMT hosts. **b,** Bar plot displaying the frequency of indicated genomic states, delineated by clustering of scRNA-seq data using ICGS2 reference clusters and cellHarmony **(Methods)**. **c,** ELISPOT analysis of NP-specific IgG1^+^ ASCs detected in indicated subsets measured at 35 d.p.i.. **d,** Titers of NP-specific IgG1 antibodies in µMT recipients following adoptive transfer of CD138^+^ splenic subsets (35 d.p.i.) measured at 21 d.p.t.. **e,** ELISPOT analysis of NP-specific IgG1^+^ BMPCs detected in µMT recipients following adoptive transfer of indicated CD138^+^ subsets (60 d.p.t.). Each symbol represents an individual mouse (d,e). Data are pooled from three different experiments (c,d,e). Statistical significance was tested by Kruskal Wallis with Dunn’s multiple comparison test (d,e). *p<0.05; and **p<0.01.

To complement the adoptive transfer system and also further resolve genomic states of splenic PC precursors that give rise to BMPCs, we performed clonal and genomic analyses of antigen-specific cells within the spleen and bone marrow of individual NP-KLH immunized mice. Importantly, this experimental design enabled the tracing of clones of antigen-specific cells and their genomic states in the spleen and bone marrow, in the context of an ongoing immune response, without requiring cell isolation and adoptive transfer. For such analyses, indicated CD138^+^ splenic and bone marrow subsets, with PC precursor activity (Fig. 4e), were isolated from individual mice immunized with NP-KLH (35 d.p.i.) and analyzed by BCR-seq and scRNA-seq (Fig. 5a). Consistent with the results from the adoptive transfer model system, this analysis revealed that PC-*Tigit*\*, PC-*Slpi*\* as well as PC-*Lag3*\* antigen-specific cells in the spleen could be linked to clonal derivatives manifesting corresponding genomic states in the bone marrow (Fig. 5b and Supplementary Table 5). Quantitative analysis of the clonal trajectories and distributions in the spleen and bone marrow compartments revealed a gradient of PC precursor activity in the GC response (35 d.p.i.), reflected as PC-*Tigit*\* > PC-*Slpi*\* > PC-*Lag3*\* (Fig. 5c). Furthermore, within each splenic PC precursor state, the clonal progeny in the bone marrow were predominantly reflective of that state. This experiment was repeated by including analysis of the splenic PC progenitors alongwith the various PC clusters. As noted earlier (Fig. 3c,d) we were able to detect clonal progeny of PC progenitors in the spleen but not in the bone marrow (Extended Data Fig. 5b,c and Supplementary Table 5). Collectively, these results suggest that antigen-specific, *Tigit*-expressing splenic PCs, generated in the context of a GC-dependent immune response represent the dominant source of BMPC precursors.

**Fig. 5.**
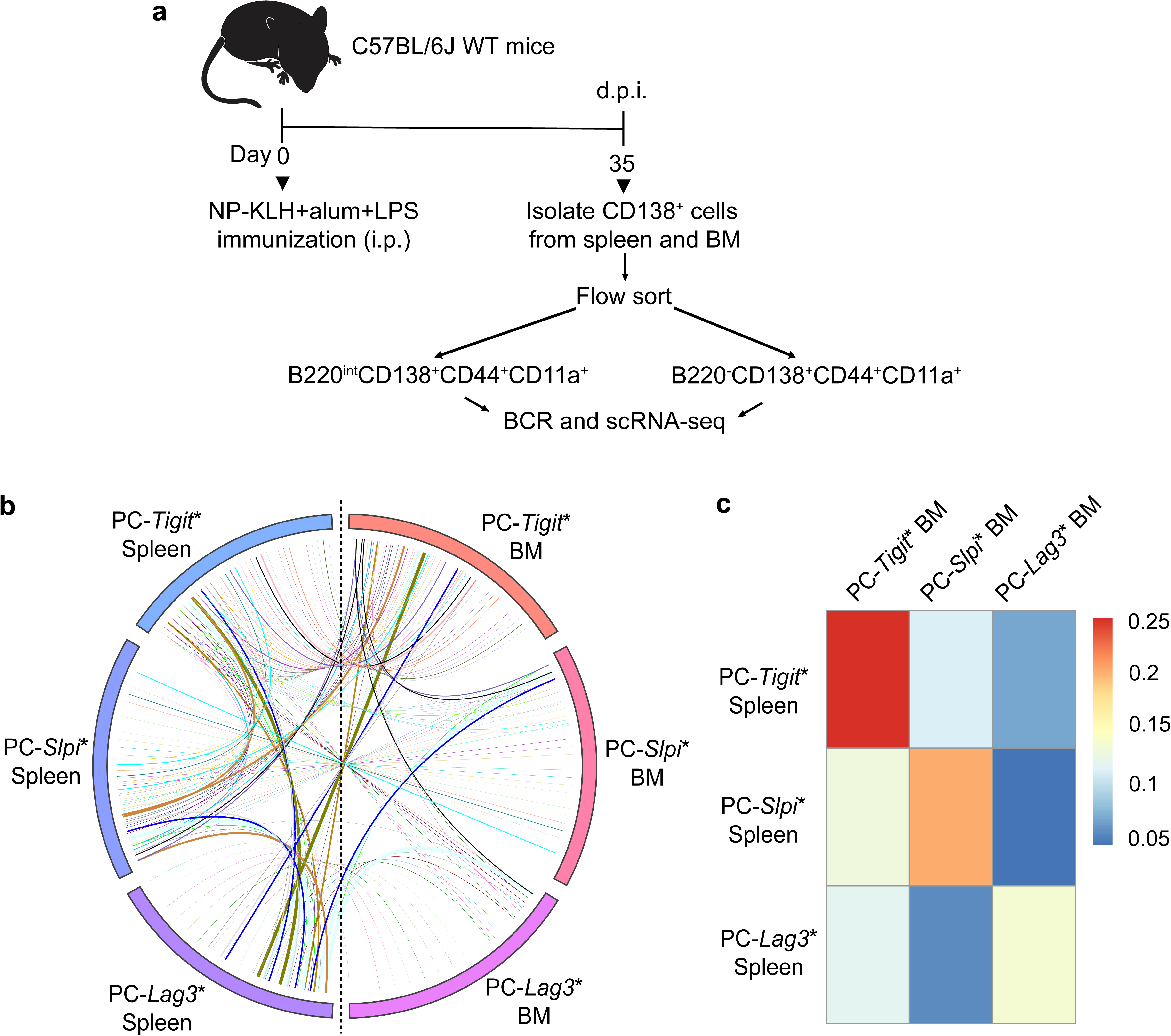
Clonal tracking of antigen-specific PC precursors migrating from spleen to bone marrow. **a,** Experimental design enabling clonal tracking of PC precursors that migrate from spleen to bone marrow of NP-KLH immunized mice (35 d.p.i.). Coupled scRNA-seq and BCR-seq was performed on indicated cells, within each compartment, isolated by flow cytometry. **b,** Circos plot displaying clones and their genomic states in spleen and bone marrow. Colored bars denote distinctive ICGS2 delineated genomic states in spleen and bone marrow. Colored lines represent clones that contain cells with identical V(D)J rearrangements that span two or more genomic states. **c,** Heatmap displaying the frequencies of clones spanning indicated genomic states in the spleen and bone marrow.

### TIGIT promotes PC precursor expansion and generation of PCs

The above results implied a cell-intrinsic function of TIGIT in the generation and/or proliferation of PC precursors. In line with the expression dynamics of the *Tigit* gene in the scRNA-seq datasets, TIGIT protein was highly expressed in splenic B220^int^CD138^+^ cells, with reduced levels in B220^−^CD138^+^ cells (Extended Data Fig. 6a). Of note, naïve and GC B cells did not express TIGIT. To analyze the cell-intrinsic functions of TIGIT, we generated mixed bone-marrow chimeras by transferring cells from CD45.1 *Tigit*^+/+^ (WT) and CD45.2 *Tigit*^−/−^ mice at a 50:50 ratio into irradiated WT recipients (Extended data Fig. 6b). CD45.2 *Tigit*^+/+^ bone marrow cells were also co-transferred with CD45.1 *Tigit*^+/+^ cells to generate control chimeras. Successful chimeras were confirmed by a 1:1 ratio of CD45.1 WT and CD45.2 *Tigit*^−/−^ naive B cell populations in the spleen at 8 weeks after transplantation (Extended Data Fig. 6c), indicating that the deficiency of TIGIT did not impact the overall development of B cells. Notably, the proportions of *Tigit^−/−^* cells were significantly decreased in the B220^int^ and B220^−^ PC compartments in the spleen and bone marrow (Extended Data Fig. 6d,e), demonstrating that TIGIT was required in a cell-intrinsic manner to maintain the steady-state frequencies of plasma cells and their precursors. To analyze the function of TIGIT in the generation of antigen-specific plasma cells, we immunized chimeric mice with NP-KLH and analyzed the wild-type and *Tigit^−/−^* splenic PC compartments at the indicated time points (Fig. 6a). Consistent with the observations in chimeric mice at steady state, the proportions of *Tigit*^−/−^ cells in the splenic plasma cell precursor compartment were significantly decreased after immunization as were their differentiated PC progeny (Fig. 6b-d). Importantly, the numbers of NP-specific ASCs generated from the *Tigit*^−/−^ B cells were also significantly decreased in the spleen and bone marrow of the chimeras (Fig. 6e,f). Thus TIGIT functions in a cell-intrinsic manner to promote the generation and/or proliferation of PC precursors. Given that TIGIT-expressing PC precursors appear to undergo greater clonal expansion (Fig. 3f), we analyzed their proliferation index based on KI-67 staining (Extended Data Fig. 6f,g). Notably, a larger fraction of TIGIT^+^ cells within the B220^int^CD138^+^CD44^+^CD11a^+^ PC precursor compartment were KI-67^+^ suggesting that TIGIT could function in controlling the proliferation of PC precursors. We tested this possibility by performing EdU incorporation assays *in vivo* with the immunized chimeric mice. The deficiency of TIGIT significantly impaired the proliferation of plasma cell precursors (Fig. 6g,h). Thus, TIGIT-mediated signaling in GC-derived PC precursors controls their expansion and the generation of BMPCs.

**Fig. 6.**
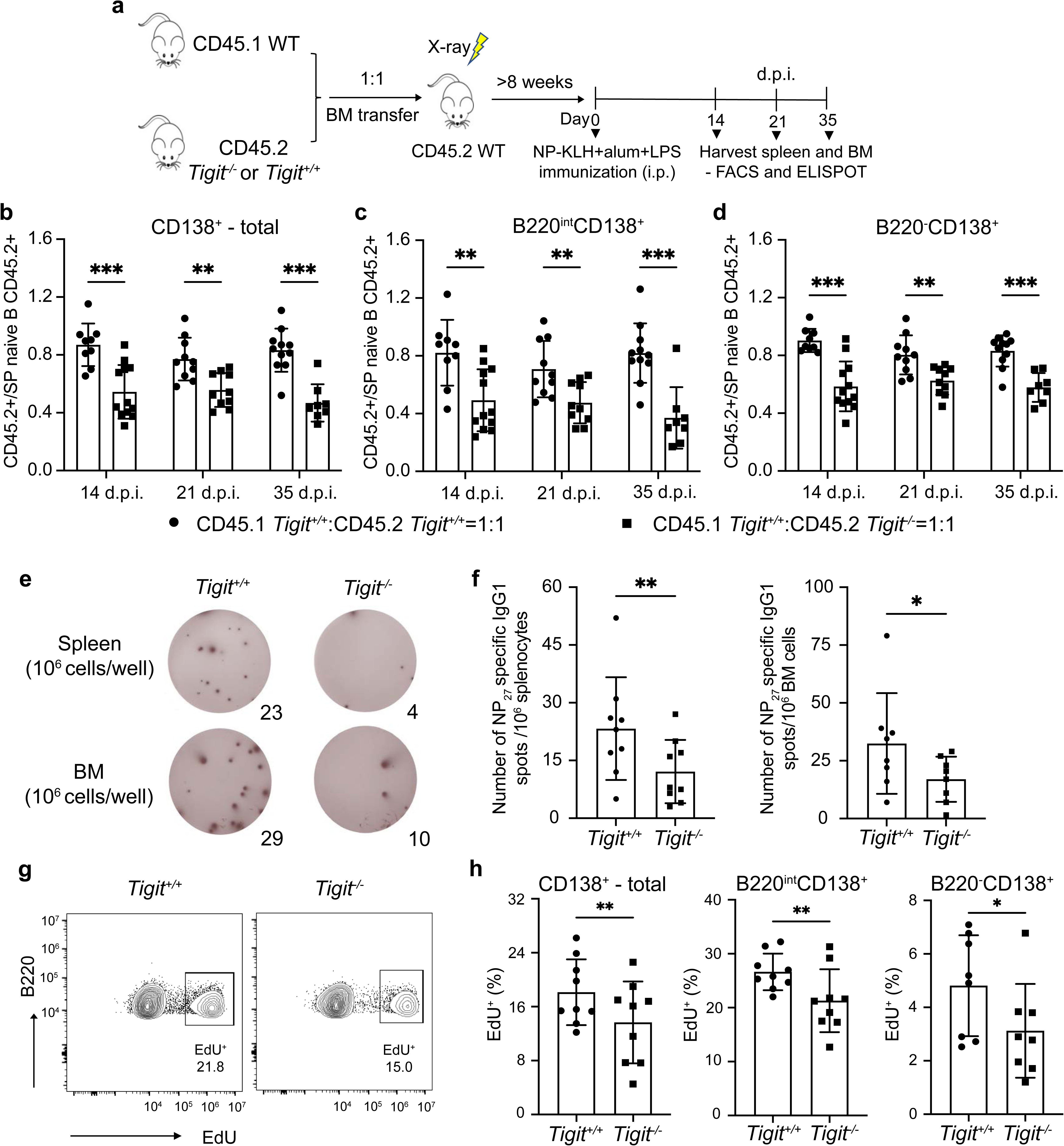
*Tigit* deficiency impairs generation of PCs. **a,** Experimental design of the bone marrow chimeric mouse model used to analyze function of *Tigit* in the generation of PCs after NP-KLH immunization. **b-d**, Quantitative analysis of the normalized proportions of CD45.2^+^ cells in the CD138^+^ splenic compartment of control (CD45.1 *Tigit^+/+^*:CD45.2 *Tigit^+/+^=*1:1) and experimental (CD45.1 *Tigit^+/+^*:CD45.2 *Tigit^−/−^=*1:1) chimeric mice: **(b)** total, **(c)** B220^int^ and **(d)** B220^−^ subsets at the indicated d.p.i.. The proportions of CD45.2^+^ cells in the various CD138^+^ subsets were normalized based on the fraction of CD45.2^+^ naïve splenic B cells in each chimeric mouse. **e,f,** ELISPOT analysis of NP-specific IgG1^+^ ASCs (*Tigit^+/+^* or *Tigit^−/−^*) in spleen and bone marrow compartment of chimeric mice after NP-KLH immunization (35 d.p.i.). Splenic and bone marrow cells were sorted based on CD45.1 or CD45.2 expression and plated for ELISPOT analyses. Representative images of NP-specific IgG1^+^ ASCs at 35 d.p.i. **(e)** and quantitative analysis **(f)**. **g,** Representative flow plots showing the percentage of EdU^+^ cells in B220^int^CD138^+^ subsets (*Tigit^+/+^* or *Tigit^−/−^*) in the spleen of chimeric mice after NP-KLH immunization (35 d.p.i.). **h,** Plots displaying the EdU^+^ cells in total (left), B220^int^ (middle) and B220^−^ (right) CD138^+^ subsets (*Tigit^+/+^* or *Tigit^−/−^*) in the spleen of chimeric mice after NP-KLH immunization (35 d.p.i.). Each symbol represents an individual mouse (b-d,f,h). Data are representative of three independent experiments (b-h) and shown as the mean ± SEM. Statistical significance was tested by two-tailed t-test (b-d,f,h). *p<0.05; **p<0.01; ***p<0.005 and ****p<0.0001.

## Discussion

We have attempted to address several fundamental questions concerning the nature of PC precursors that emanate from a secondary lymphoid organ, via a GC B cell response, and migrate to the bone marrow to give rise to PCs of varying longevity. Using the adoptive transfer of CD138^+^ splenocytes, isolated at varying times during an NP-KLH immunization response, into naïve B cell-deficient mice, we demonstrate a quantitative as well as qualitative change in PC precursor activity. Strikingly, the peak of PC precursor activity is observed at 35 d.p.i., 3 weeks after the peak of the GC response (14 d.p.i.). Furthermore, such precursors preferentially give rise to longer-lived PCs in the bone marrow. Consistent with our findings, temporal analysis of NP-specific ASCs in spleen and BM, during a NP-CGG response, suggested a late (29-47 d.p.i.) emergence of long-lived PCs that coincided with their maximum output in the splenic compartment^12,13^. It should be noted that the two experimental systems generate concordant results despite the substantial differences in approaches. The former involves time-resolved and quantitative analysis of newly generated NP-specific ASCs in spleen and bone marrow compartments of immunized animals, but did not involve the characterization and functional testing of PC precursors. Our approach, involving the adoptive transfer of PC precursors isolated from immunized animals into naïve B cell-deficient hosts, also generates their temporal and quantitative activity profiles. It additionally enables molecular and functional characterization of PC precursors, in particular the linking of genomic states with capacity to home to the bone marrow as well as assessment of longevity. Recently, a distinctive experimental approach, using a PC time-stamp inducible Cre-recombinase mouse model controlled by *Prdm1* demonstrated that LLPCs can begin to accumulate in the BM starting as early as 3 d.p.i. in an immune response^39^. We note that in the time-stamp model, LLPCs were tracked up to 70 days. We demonstrate using our adoptive transfer system that PC precursors generated at 21 d.p.i. can give rise to PCs that are detectable between 50-80 total days, after immunization. Notably, by analyzing PC precursors that arise much later in the GC response at 35 d.p.i. we show that they give rise to longer lived PCs in the bone marrow, up to a total of 155 days after immunization. The collective evidence is consistent with both quantitative and qualitative changes in the PC precursor compartment, as the germinal center response wanes due to diminished antigen availability.

Though the adoptive transfer experiments of CD138^+^ splenocytes enabled temporal and quantitative monitoring of PC precursor activity, they did not reveal the identity of the cells nor their genomic states. To do so unequivocally, we performed scRNA-seq and BCR-seq of CD138^+^ splenocytes and subsets so as to link their genomic states with developmental capacities. The conclusions of these experiments were then validated in immunized but otherwise unperturbed mice, by clonal tracing of antigen-specific PC precursors that were generated in the spleen and gave rise to PCs in the bone marrow. Genomic analysis of the CD138^+^ splenocyte compartment revealed a novel transition state that we have termed PC progenitors. These transitional cells appear to be inducing the transcription factors IRF4, BLIMP1 and XBP1 and consequently the expression of a large number of PC genes, while retaining the expression of B cell identity genes and down-regulating BCL6 and other GC genes. This suggests that the repression of B cell identity genes which is a hallmark of plasma cells occurs subsequent to the specification of the PC fate. It remains possible that the gene activation and repression mechanisms are concomitantly deployed in PC progenitors but the latter require slower acting chromatin modifications and therefore manifest with delayed kinetics in their transcript dynamics. Consistent with the genomic analysis of PC progenitors the cells reside within the B220^+^ subset of CD138^+^CD44^+^CD11a^+^ splenocytes. Notably they secrete antigen-specific antibodies, have acquired somatic hypermutations and appear to be undergoing a G1 to S phase transition based on DEG analysis of PC progenitors at 35 d.p.i. vs. 21 d.p.i.. This analysis suggests that the G1 to S phase cell cycle transition occurs in a more pronounced fashion at 35 d.p.i. which is in keeping with the peak of PC precursor activity revealed by our adoptive transfer experiments.

The foregoing analysis led us to consider the possibility that the PC progenitors, may represent the sought after PC precursors, whose activity is temporally and quantitatively monitored in the adoptive transfer experiments. However two complementary sets of experimental approaches ruled out this possibility and instead demonstrated that the dominant set of PC precursors are differentiated progeny of the PC progenitors. The majority of the PC precursors reside within a splenic PC cluster that express, in an inducible manner, the inhibitory T cell immunoreceptor TIGIT. This conclusion is based on two key findings: (i) that the B220^+^ subset of CD138^+^CD44^+^CD11a^+^ splenocytes within which PC progenitors reside have undetectable PC precursor activity in the adoptive transfer experiments; in contrast, the B220^int^ subset of CD138^+^CD44^+^CD11a^+^ splenocytes, containing the majority of PC-*Tigit\** cells, have appreciable PC precursor activity; and (ii) clonal tracing demonstrates that splenic PC-*Tigit\** cells are the predominant source of PC lineages in the bone marrow. We note that in the adoptive transfer experiments, we were not able to quantitatively distinguish between the PC precursor activity of the three PC clusters revealed by the scRNA-seq analysis, namely the PC-*Tigit\**, PC-*Slpi\** and PC-*Lag3\** cells. However the clonal tracing experiments, not involving cell transfers, enumerated the PC-*Tigit\** cells as the major source of PC precursors that give rise to heterogeneous PCs in the bone marrow. Notably, the PC-*Tigit\** cells have increased expression of a mitotic gene module, manifest increased EdU labeling, and undergo greater clonal expansion than their PC progenitor counterparts. Consistent with these inferences drawn from the single-cell analyses, TIGIT functions in a cell-intrinsic manner to promote the proliferation of splenic PC precursors and the consequent generation of PCs in the bone marrow (discussed below). Complimenting our findings, a recent analysis of antigen-specific splenic and bone marrow PC compartments using scRNA-seq and pulse-chase experiments in mice revealed that LLPCs are heterogenous and have distinct surface phenotypes that are associated with different Ig isotypes. IgA-expressing LLPCs were Ly6A^hi^TIGIT^−^, whereas IgG and IgM-expressing LLPCs were EPCAM^hi^CXCR3^−40^. Our clonal tracing analysis suggests that PC-fate-specified cells generated from SLOs/GCs expressing *Tigit* and *Cxcr3* (35 d.p.i.) may preceed the emergence of such heterogeneous LLPCs. Thus we propose a two-step model for the generation of functionally competent PC precursors that can migrate from the spleen and home to the bone marrow to undergo terminal differentiation. The first step involves the generation of a PC progenitor, that manifests a mixed genomic state, reflected by induction of an initial program of PC genes while retaining expression of B cell identity genes. PC progenitors appear to be exiting from the GC based on downregulation of GC-specific genes and their single-cell trajectories. The PC progenitor trajectories inferred by two complementary computational methods, Monocle and scVelo, are continuous paths that merge with PC clusters. Notably, PC progenitors are not functionally competent as PC precursors as they may not have upregulated the expression of proteins needed for cell migration and homing to long-lived niches such as the bone marrow or gut. Such cellular programming is proposed to comprise the second step and converts the PC progenitor into a PC precursor with migratory and niche homing functions. Importantly, the conversion of the PC progenitor into a PC precursor is accompanied with the robust expression of the PC gene program, downregulation of B cell identity genes and proliferative expansion that is dependent on the inducible expression of the inhibitory signaling receptor, TIGIT.

Three dominant signaling systems that function to regulate the dynamics of B cells during a GC response involve the BCR, CD40 and IL-21R. BCR and CD40 signaling, the latter enabled by interaction with cognate Tfh cells, are essential for the positive selection of GC B cells and their cycling in the LZ and DZ regions^41,42^. Conversely, IL-21 signaling in conjunction with CD40 appears to promote the exit of B cells from the GC and the generation of PC precursors^24^. In keeping with these findings, NP-reactive B cells isolated at the peak of GC response, when stimulated with CD40L and IL-21 for 48 hours, generated BCL6^lo^IRF4^hi^ cells^25^ that likely represent PC precursors. However, if BCR signaling was stimulated along with CD40 and IL-21R, the GC B cells preferentially generated precursors of memory B cells^25^. In keeping with these findings, the IL-21 promoted differentiation of human tonsillar B cells into plasma cells was inhibited by BCR signaling^43^. We propose that BCR signaling antagonizes IL-21R signaling, and therefore the number of PC precursors that are generated via a GC response would be predicted to be inversely proportional to antigen availability *in vivo*. As antigen availability and therefore BCR signaling are diminished during the later phases of a GC response, enhanced IL-21 signaling in conjunction with CD40 would synergize to increase generation of PC precursors. We note that loss of IL-21R signaling has been also shown to impact the number of antigen-specific ASCs and antibody titers at late stages in NP- and viral antigen-responses^44^. These results raise the possibility that the magnitude and/or duration of IL-21 signaling in GC B cells, possibly as a consequence of their repeated antigen and Tfh encounters, could induce differentially programmed PC precursors that give rise to longer-lived PCs.

The model, invoking the antagonistic interplay between BCR and IL-21 signaling in the generation of PC precursors, raises a paradoxical role for TIGIT signaling, which we show to be important for the proliferation and expansion of PC precursors. TIGIT is a coinhibitory receptor, containing an ITIM signaling domain, expressed on the surface of subsets of T, NK and B cells^45^. TIGIT ligands include the poliovirus receptor (PVR) and Nectin-2 (PVRL2)^45^. Loss of TIGIT in T lineage cells results in susceptibility to various autoimmune diseases^45^. There is limited evidence for the functioning of TIGIT in B cells. It is expressed on the surface of a subset of regulatory B cells and its deletion in the B-lineage results in the development of CNS inflammation and spontaneous paralysis in mice^46,47^. TIGIT inhibits TCR signaling and its loss or antagonism promotes T cell hyperproliferation^48^. We demonstrate that TIGIT is expressed in an inducible manner in PC precursors and functions in a cell-intrinsic manner to promote their proliferation. These seemingly paradoxical findings can be reconciled in the context of our model that invokes antagonistic interplay between BCR and IL-21 signaling in the generation and expansion of PC precursors. We propose that TIGIT inhibits BCR signaling in a manner analogous with its functions in antagonizing TCR signaling, thereby enabling the generation and proliferation of PC precursors promoted by IL-21 in conjunction with CD40 signaling. In the model, IL-21 and CD40 signaling programs GC B cells to exit the germinal center and also to undergo clonal expansion. This aspect is analogous to the role of BCR and CD40 signaling in programming GC B cells to migrate to the dark zone and undergo mitotic divisions. The direct demonstration of an inhibitory function of TIGIT in BCR signaling, the nature and expression of its ligand and the underlying molecular mechanisms remain to be elucidated. It remains possible that TIGIT antagonizes signaling by a distinct receptor expressed in PC precursors. Overall we propose that the interplay of BCR, CD40 and IL-21 signaling early in the GC response when antigen is not limiting favors BCR and CD40 signaling thereby reducing the generation of PC progenitors and precursors that exit the GC. As antigen becomes limiting, later in the GC response, BCR signaling is diminished enabling persistent IL-21 signaling in conjunction with CD40 to increase the frequency of PC progenitors and precursors. The inducible expression of TIGIT initiated in PC progenitors, potentially due to repeated antigen encounters by GC B cells, could amplify the generation of PC precursors by further impairing BCR signaling thereby promoting IL-21 signaling. These dynamic alterations in the interplay of the BCR, TIGIT, CD40 and IL-21 signaling systems in the context of an ongoing GC B cell response could not only impact the numbers of PC precursors that are generated at early versus later stages of the response but also their genomic programming and therefore the longevity of the PCs that arise in the bone marrow. In keeping with this suggestion analysis of DEGs in the PC-*Tigit\** cells (35 d.p.i. vs. 21 d.p.i.) revealed that PC precursors arising later in the GC response exhibit higher expression of genes involved in focal adhesion and ribosomal genes. Coordinated expression of both sets of genes have been shown to be important for cellular interactions with the extracellular matrix by promoting localized translation of focal adhesion proteins^49–51^. Thus, we propose a model for the proliferation and programming of precursors of long-lived PCs, based on extended antigen encounters followed by reduced antigen availability.

The above model has important implications for durable antibody responses to pathogens. It implies that robust extrafollicular B cell responses which clear microbial antigens rapidly would generate lower affinity and shorter-lived PCs. In contrast, extended GC responses, driven by continued antigen availability, would not only favor the generation of higher affinity but longer lived PCs. The model can be tested in the context of pathogen as well as vaccine responses. Importantly, our model posits the coordination of affinity maturation occurring via somatic hypermutation and positive selection in the germinal center with the differential genomic programming of PC precursors that exit the GC and home to the bone marrow. It makes key predictions regarding the interplay of BCR, TIGIT and IL-21 signaling in the generation of longer lived PCs that will be experimentally tested in future experiments. The temporally resolved scRNA-seq datasets of NP-specific B cell clones in the current manuscript along with those in our earlier studies^21,31^ represent a unique and extensive catalog of genes. Many of these genes likely function in the generation of PC precursors, their migration and homing as well as in niche-specific interactions of PCs. (http://www.altanalyze.org/ICGS/Public/Splenic_and_Bone_Marrow_Plasma_Cell_Compartments/User.php). The adoptive transfer approach used by us is generalizable and could be extended to analyze PC precursors that home to other anatomical locations and give rise to PCs in diverse niches.

---

**Extended Data Fig. 1.**
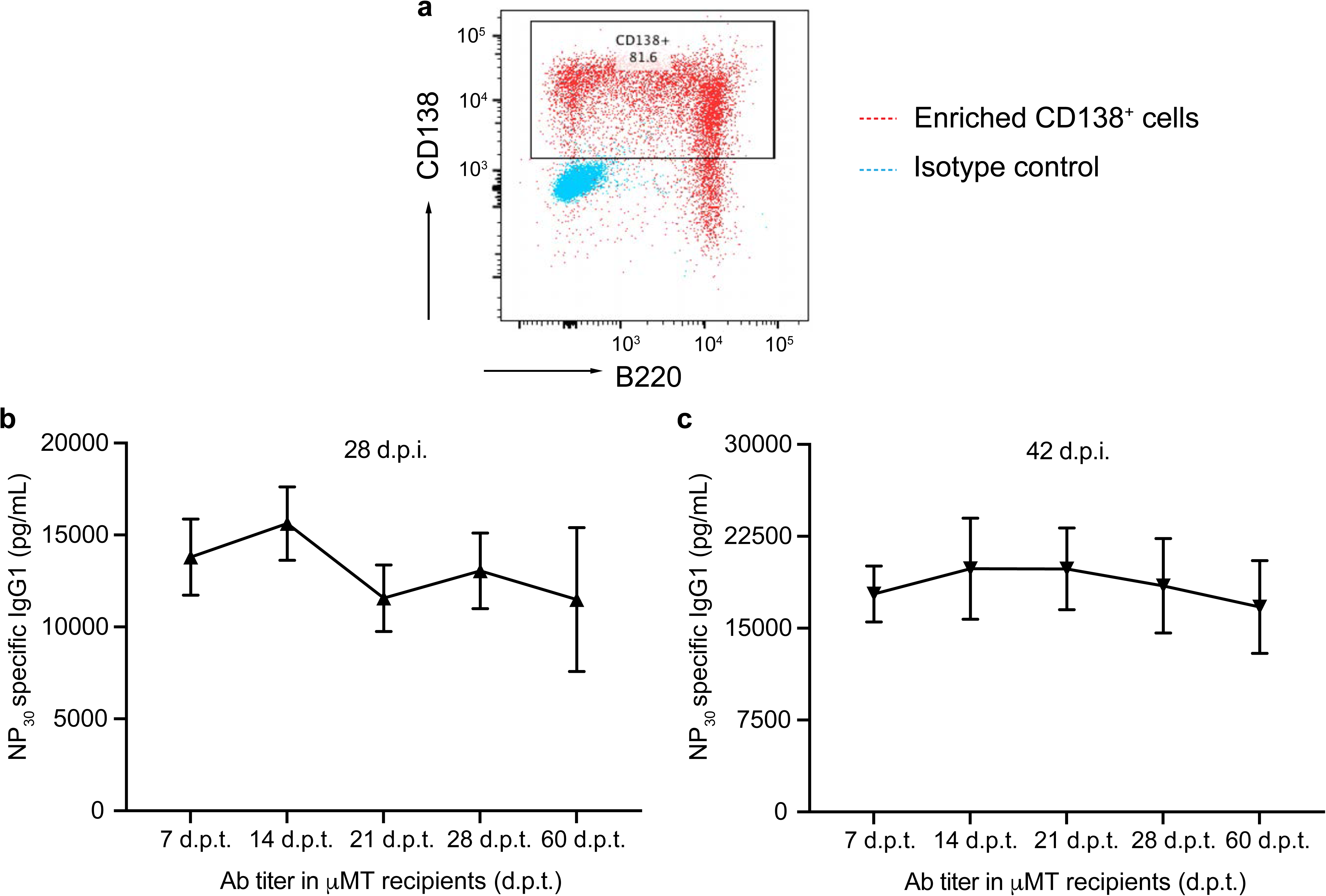
Temporal dynamics of BMPC precursors generated during a GC response. **a,** A representative flow plot showing the frequency of CD138 expressing cells in the enriched CD138^+^ splenocytes (positive selection) that were used in adoptive transfer experiments. **b,c,** Titers of NP-specific IgG1 antibodies in µMT recipients following adoptive transfer of CD138^+^ splenocytes at 28 d.p.i. (n=6) **(b)** and 42 d.p.i. (n=7) **(c)** at indicated d.p.t..

**Extended Data Fig. 2.**
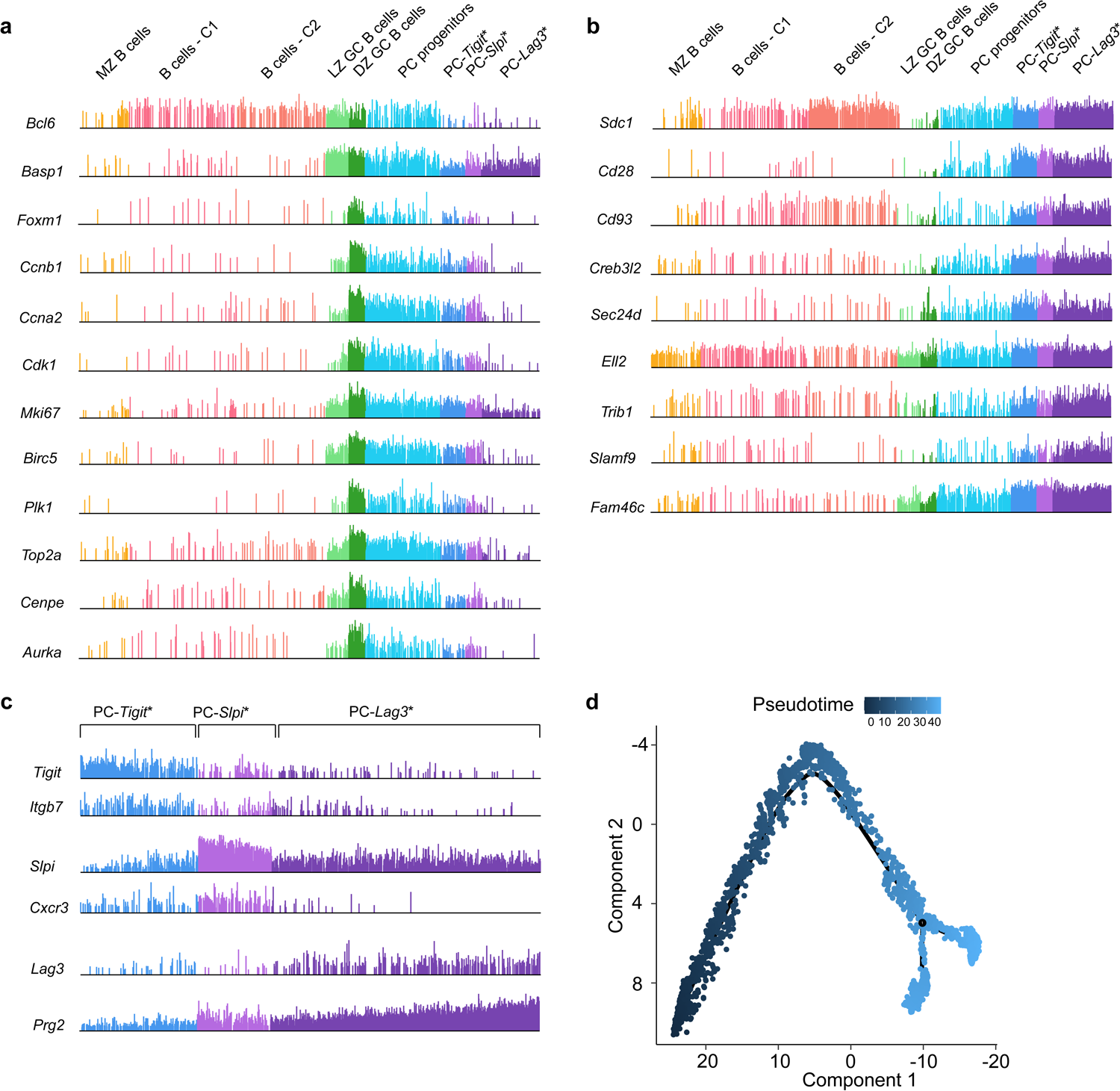
scRNA-seq analysis of PC genomic states and trajectories. **a,** Comb plots displaying the incidence and amplitude of indicated GC B cell, DZ GC and cell cycle genes in designated cluster as in Fig. 2a. **b,** Plots displaying the incidence and amplitude of indicated PC genes in each cluster as above. **c,** Plots displaying the incidence and amplitude of indicated marker genes distinguishing the three PC clusters as above. **d,** Pseudotime analysis of cells in the Monocle 2 trajectory (Fig. 2e). LZ GC B cells were set as the root state for the pseudo trajectory analysis.

**Extended Data Fig. 3.**
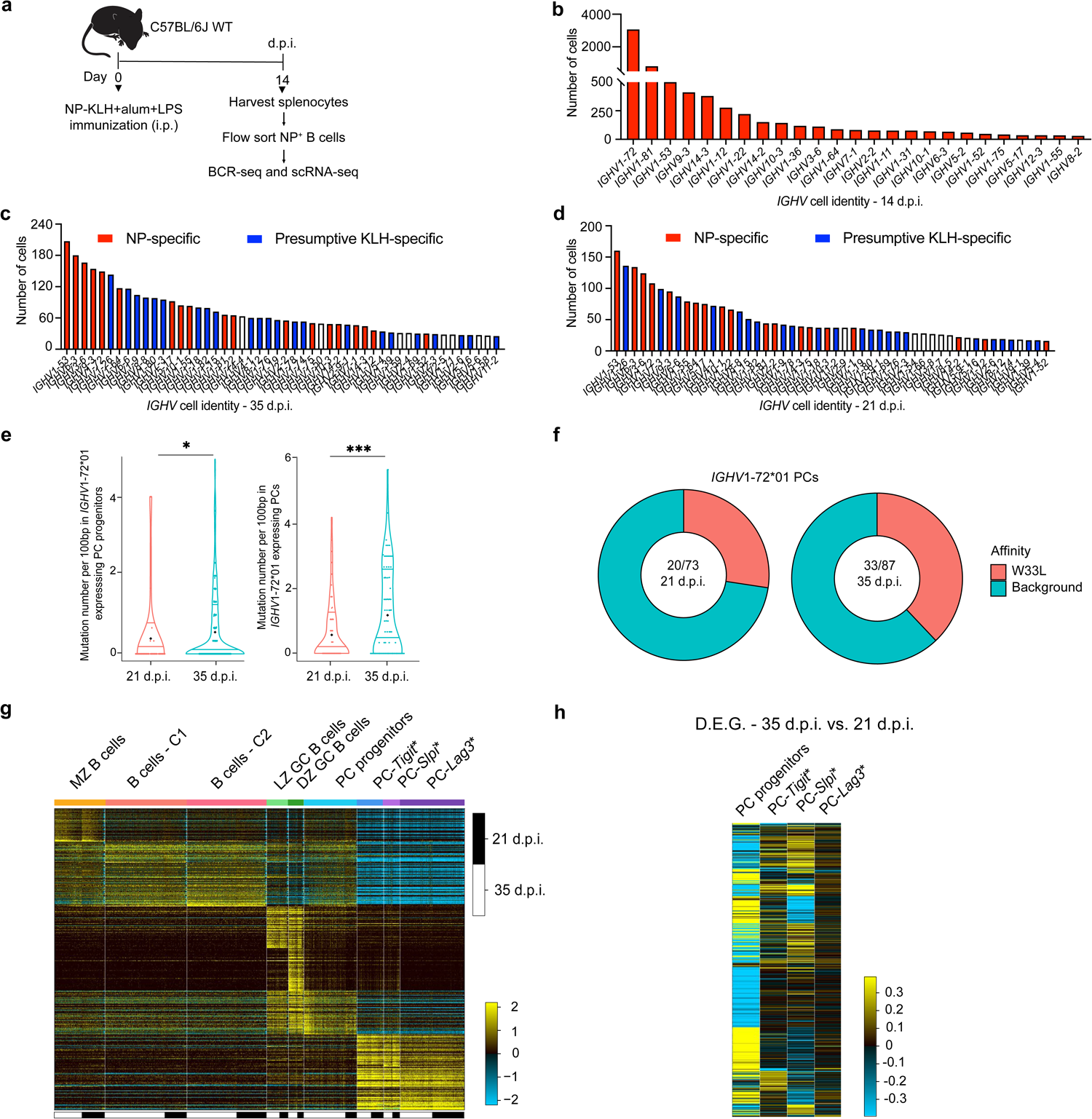
Delineation of antigen-specific PC genomic states and clonal tracking. **a,** Experimental design for isolation and analysis of NP-specific B220^+^ cells from C57BL6/J mice (14 d.p.i.), analyzed by scRNA-seq and BCR-seq. **b,** Plot displaying the rank order of top 25 *IGHV* genes expressed in NP-specific B220^+^ cells (14 d.p.i.). **c, d,** Plots displaying the rank order of top 50 *IGHV* genes expressed in CD138^+^ splenocytes on 35 d.p.i. **(c)** and 21 d.p.i. **(d).** Red bars are NP-specific *IGHV* genes expressed on 14, 21 and 35 d.p.i., blue bars are presumptive KLH-specific *IGHV* genes expressed on 21 and 35 d.p.i.. **e,** Mutation frequencies of *IGHV1-72*01* gene in splenic PC progenitors (left) and PC clusters (right) on 21 and 35 d.p.i.. **f,** Pie-charts displaying the proportion of *IGHV*1-72*01 gene segments harboring high affinity W33L mutations in splenic PCs (21 d.p.i. vs. 35 d.p.i.). **g,** Combined heatmap of genomic states in the enriched CD138^+^ splenocytes from both 21 and 35 d.p.i., generated using cellHarmony, for which CD138^+^ splenocytes (35 d.p.i.) were used as a reference dataset. Columns in heatmap represent cells from 21 and 35 d.p.i. (n=14,068); rows represent MarkerFinder genes (n=413). **h,** Heatmap of differentially expressed genes (DEGs) in NP- and KLH-specific PC progenitors and indicated PC genomic states (35 vs. 21 d.p.i.) generated using cellHarmony, for which 35 d.p.i. were used as a reference dataset. Rows represent the expression of differentially expressed genes on 35 d.p.i.. Statistical significance was tested by Wilcoxon test (e). *p<0.05; and ***p<0.005.

**Extended Data Fig. 4.**
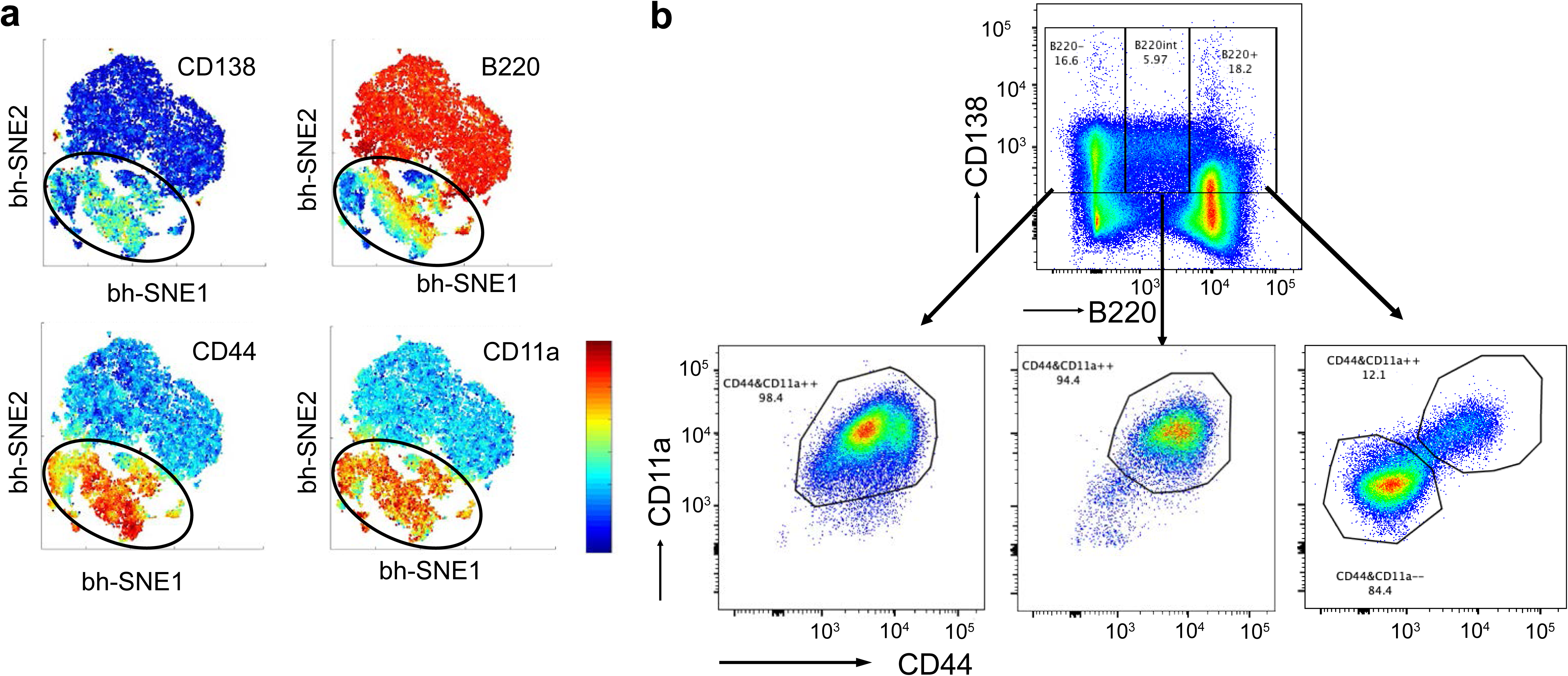
Functional and genomic analysis of BMPC precursors within splenic CD138^+^ subsets. **a,** Representative t-SNE plots based on flow cytometry data showing the indicated markers in the enriched CD138^+^ splenocytes obtained using positive selection at 35 d.p.i.. **b,** Representative flow plot showing the gating strategy for sorting CD138^+^ splenic subsets (35 d.p.i.) used in adoptive transfer experiments. Purity of the indicated PC subsets are displayed.

**Extended Data Fig. 5.**
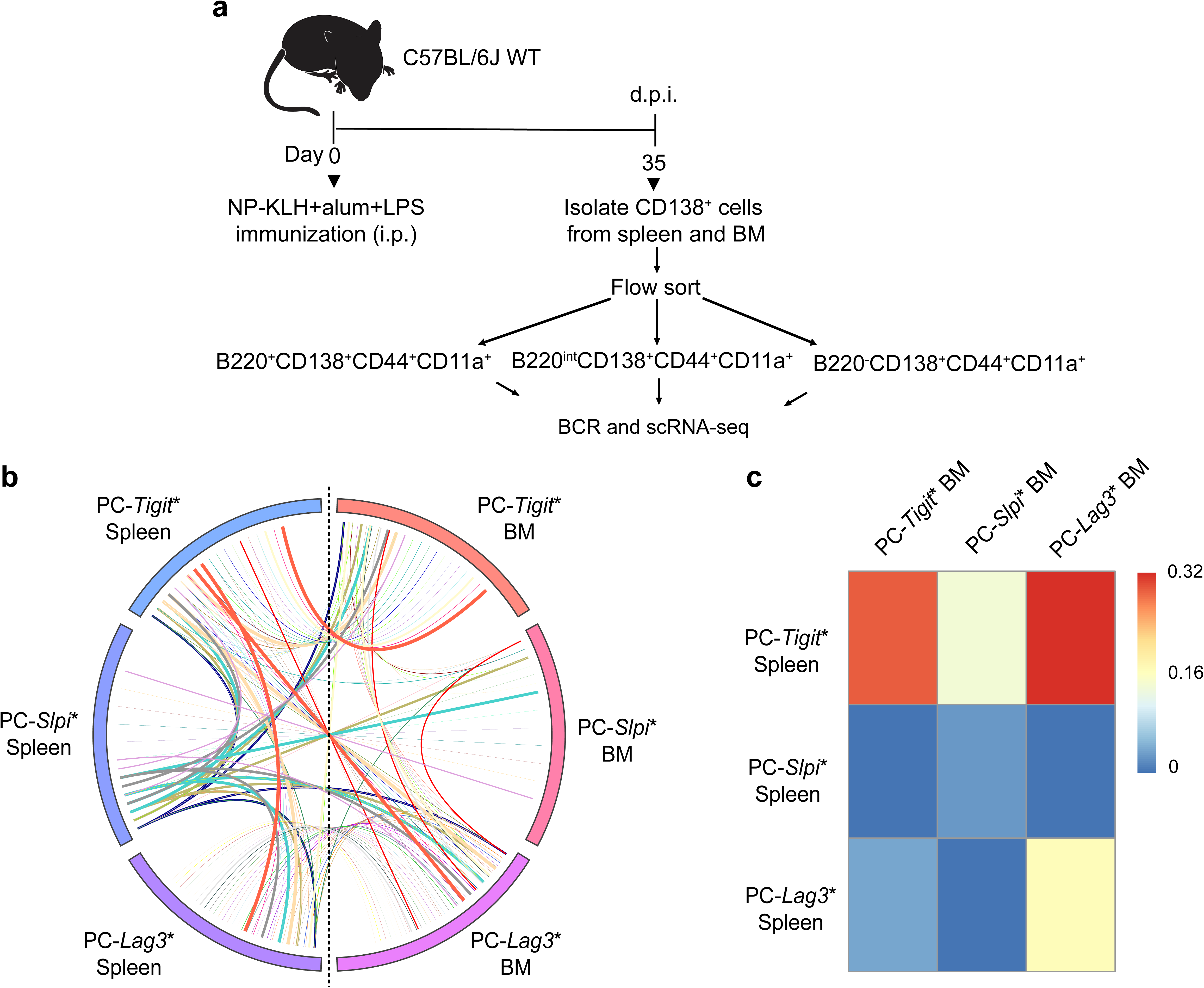
Clonal tracking of antigen-specific PC precursors that migrate from spleen to bone marrow. **a,** Experimental design enabling clonal tracking of PC precursors that migrate from spleen to bone marrow of NP-KLH immunized mice (35 d.p.i.). Coupled scRNA-seq and BCR-seq was performed on indicated cells (including PC progenitors), within each compartment, isolated by flow cytometry. **b,** Circos plot displaying clones and their genomic states in spleen and bone marrow. Colored bars denote distinctive ICGS2 delineated genomic states in spleen and bone marrow. Colored lines represent clones that contain cells with identical V(D)J rearrangements that span two or more genomic states. **c,** Heatmap displaying the frequencies of clones spanning indicated genomic states in the spleen and the bone marrow.

**Extended Data Fig. 6.**
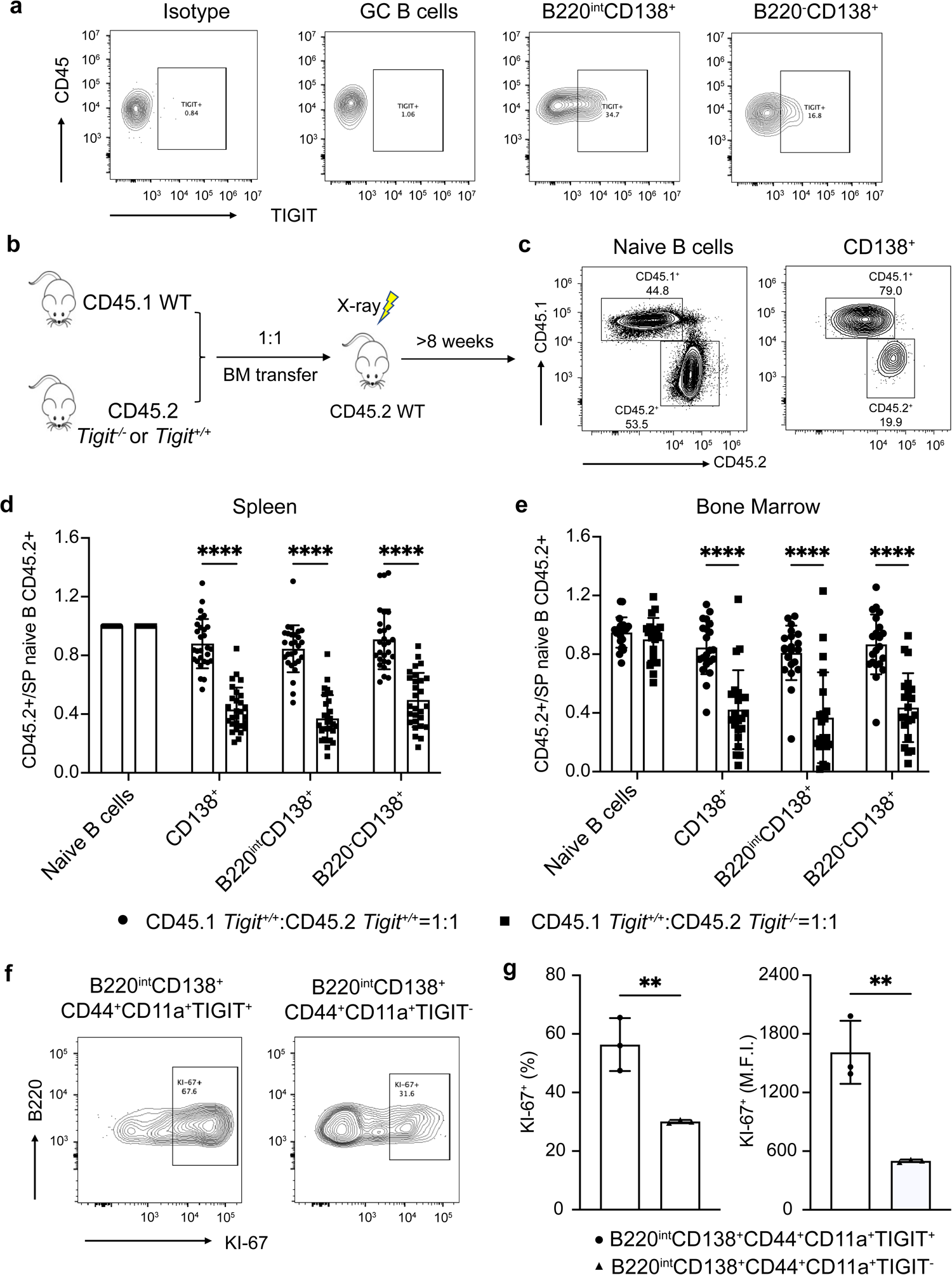
***Tigit* deficiency impairs generation of PCs a**, Representative flow plots showing TIGIT expression in B cell and PC subsets. **b**, Experimental design for the bone marrow chimeric model. **c**, Representative flow plots showing the proportions of CD45.1^+^ or CD45.2^+^ cells in each cell compartment of the chimeras. **d, e,** Quantification of the normalized proportions of CD45.2^+^ cells in the indicated cell compartments in the spleen **(d)** and bone marrow **(e)** of the chimeras. The proportions of CD45.2^+^ cells in individual compartments were normalized to the proportion of CD45.2^+^ cells within naïve B cell subset in the spleen of each mouse. **f**, Representative flow plots showing the frequencies of KI-67^+^ cells in TIGIT-expressing cells and their counterparts in the B220^int^ plasma cell subset (35 d.p.i.). **g**, Quantification of the frequency and mean fluorescence intensity of KI-67^+^ cells in B220^int^ subset as in **(f)**. Each symbol represents an individual mouse (d,e,g). Statistical significance was tested by two-tailed t-test (d,e,g). **p<0.01; and ****p<0.0001.

## Methods

### Mice

C57BL/6J (CD45.2) (Jax 000664) and μMT (Jax 002288) mouse strains were obtained from the Jackson Laboratory. C57BL/6J and μMT mice were housed in specific pathogen-free (SPF) conditions and were used and maintained in accordance with the Institutional Animal Care and Use Committee (IACUC) guidelines in Cincinnati Children’s Hospital and Medical Center (CCHMC) and University of Pittsburgh, USA under protocol no. 2016-0096 and 19115454. *Tigit^−/−^* mice were generously provided by Dr. ZuSen Fan (Chinese Academy of Sciences). CD45.1 (Jax 002014) mice were obtained from the Jackson Laboratory. *Tigit^−/−^* and CD45.1 mice were housed in SPF conditions and were used and maintained in accordance with the IACUC guidelines of Westlake University, China under protocol no. 18002-XHP.

### Bone marrow chimeras

CD45.2 B6 recipients were lethally irradiated by X-rays (5 Gy x 2), and then intravenously transferred with a combination of 4 × 10^6^ bone marrow (BM) cells from indicated donors, mixed according to the indicated ratios. Chimeric mice were maintained with antibiotic-containing water (0.4 g/L Sulfamethoxazole) for 4 weeks after reconstitution and then returned to regular drinking water.

### Immunization

Seven to eight week old mice were immunized intraperitoneally with 100 μg NP(23)-KLH, NP(25)-KLH or NP(27)-KLH (Biosearch Technologies) mixed with 50% (v/v) imject alum (Thermo Fisher Scientific) and 1 μg LPS (Sigma) as described previously^1,2^.

### Flow cytometry staining

Splenocytes and BM cells were washed and prepared as single-cell suspensions in MACS buffer (pH 7.4; 1x PBS without calcium and magnesium plus 2% FBS and 10 mM EDTA). Erythrocytes were depleted using ACK lysis buffer (Thermo Fisher Scientific). Non-specific antibody binding was blocked by incubating the cells with Fc-blocking antibody 2.4G2 (BD, 25 μg/ml) for 10 minutes on ice. Cells were stained for 30 minutes at 4°C with viability dye eFluor 780 (eF780) Thermo Fisher Scientific Cat# 65-0864-14 and indicated antibody cocktails. All antibodies used for flow cytometry are listed below: FITC anti-mouse CD45.1 (A20) BioLegend Cat# 110706, PE anti-mouse CD45.2 (104) BioLegend Cat# 109808, Brilliant Violet 785 anti-mouse CD45 (30-F11) BioLegend Cat# 103149, Brilliant Violet 510 anti-mouse/human CD45R/B220 (RA3-6B2) BioLegend Cat# 103248, Alexa Fluor® 647 anti-mouse CD138 (281-2) BioLegend Cat# 142526, Alexa Fluor® 700 anti-mouse CD38 (90) eBioscience™ Cat# 56-0381-82, eFluor™ 450 anti-mouse/human GL7 (GL7) eBioscience™ Cat# 48-5902-82, PerCP-eFluor 710 anti-mouse TIGIT (GIGD7) eBioscience™ Cat# 46-9501-82, PE-Dazzle Red 594 B220 (RA3-6B2) BioLegend Cat# 103258, PerCP-Cy5.5 CD44 (IM7) BioLegend Cat# 103032, FITC CD11a (M17/4) BioLegend Cat# 101106 and APC Streptavidin BioLegend Cat# 405207. Flow cytometry data were collected on Cytoflex (Beckman Coulter) or LSRII (BD Biosciences) and analyzed using FlowJo 10.7.1 software (TreeStar).

### EdU labeling

EdU (Thermo Fisher Scientific) was dissolved in sterile 1x PBS (1 mg in 100 µL) and was injected intraperitoneally. For EdU detection, cells were subjected to Click reaction using Click-it EdU Flow Cytometry Assay Kit (Thermo Fisher Scientific) according to manufacturer’s instructions, followed by antibody labeling of B cell markers.

### Isolation of CD138^+^ cells and subsets

Splenic and bone marrow cells (tibia, femur and iliac crest) were isolated from C57BL/6J mice immunized with NP-KLH, alum and LPS on 21-, 28-, 35- and 42-days post-immunization (d.p.i.). Splenocytes and BM cells were washed and prepared as single-cell suspensions in MACS buffer. Enriched CD138^+^ cells were obtained by using Easysep™ Release Mouse Biotin Positive Selection Kit. Cell concentration was adjusted to 100 × 10^6^ cells per mL and incubated with 50 μL of normal rat serum for 5 minutes. Biotinylated anti-CD138 (281.2) antibody (1.0 μg/mL) was added to the cells and incubated for 5 minutes. Next the cells were incubated (5 minutes) with Easysep™ Biotin positive selection antibody (50 μL/mL) followed by addition of Easysep™ Rapidspheres (100 μL/mL) (5 minutes). MACS buffer was added to the cocktail and an Easysep™ magnet was used to remove cells that were not bound to rapidspheres. After three washes, CD138^+^ cells were obtained by incubating with Easysep™ Release Buffer for 5 minutes and placing the cells in a magnet to release them from the Rapidspheres. Purity of enriched CD138^+^ cells was determined by flow cytometry after labeling a small fraction of the cells with viability dye eFluor 780 (eF780), APC Streptavidin and PE-Dazzle Red 594 B220 (RA3-6B2) for 30 minutes at 4°C. For the isolation of CD138^+^ subsets by flow cytometry, enriched CD138^+^ cells after Rapidspheres release, were labeled with viability dye eFluor 780 (eF780), APC-Streptavidin, PE-Dazzle Red 594 B220 (RA3-6B2), PerCP-Cy5.5 CD44 (IM7) and FITC CD11a (M17/4) for 30 minutes at 4°C and were sorted as B220^+^CD138^+^CD44^−^CD11a^−^, B220^+^CD138^+^CD44^+^CD11a^+^, B220^int^CD138^+^CD44^+^CD11a^+^ and B220^−^CD138^+^CD44^+^CD11a^+^ subsets using FACSAria II (BD) with 70 μm nozzle at 4°C. Enriched CD138^+^ splenocytes or flow sorted CD138^+^ subsets were washed and suspended with 1x PBS before adoptive transfer into μMT recipients.

### Adoptive transfer of splenic CD138^+^ cells and subsets

Carrier cells for the adoptive transfer were prepared by making a single cell suspension of spleen collected from μMT mice. For adoptive transfer of total CD138^+^ splenocytes, 3 × 10^5^ enriched CD138^+^ cells and 1 × 10^6^ μMT splenocytes were injected retro-orbitally into μMT recipients. For splenic CD138^+^ subsets transfer, 1 × 10^5^ flow sorted CD138^+^ subsets and 1 × 10^6^ μMT splenocytes were injected retro-orbitally into μMT recipients.

### Serum antibody analysis

NP-specific antibody titers were analyzed in peripheral blood samples collected from mice on indicated d.p.i. and/or days post-transfer (d.p.t.) using enzyme linked immunosorbent assay (ELISA) as described previously^1^. Serum samples collected post-immunization were loaded into plates with eight 3-fold serial dilutions (1:1000 to 1:2,187,000); whereas samples after adoptive transfer were loaded into plates with eight 3-fold serial dilutions (1:10 to 1:21,870).

### Detection of antibody secreting cells (ASCs)

ASCs in the spleen or BM were analyzed by enumerating the number of NP-specific spots using enzyme linked immune sorbent spot (ELISpot) as described previously^2^. For analysis of adoptive transfer experiments involving μMT recipients, single cell suspensions were prepared from the spleen or a pair of femur, tibia, and iliac crest bones and one million splenocytes or BM cells were seeded into each well. The ASCs were enumerated by calculating based on the number of cells isolated in the six bones from each μMT recipient. To enumerate the ASCs in various flow sorted CD138^+^ subsets, 1 × 10^3^ cells were seeded into each well. Splenic and BM cells from chimeric mice were flow sorted based on CD45.1 and CD45.2 expression and used for ELISPOTs.

### scRNA-seq and BCR-seq

Enriched CD138^+^ splenocytes or flow sorted CD138^+^ splenic and BM subsets were suspended in 5% FBS and loaded along with oil and beads containing unique barcodes onto the Chromium controller (10x Genomics) to form an emulsion. Single cells contained in oil droplets were then lysed and mRNA was captured by the beads and subjected to reverse transcription and cDNA amplification to generate libraries for sequencing, according to the manufacturer’s protocol, as described previously^2^. From the cDNA libraries, one aliquot was used for fragmentation and preparation for transcriptome profiling, and another for amplification of VDJ regions using the BCR kit. Samples were then sequenced using a NovaSeq 6000 (Illumina) in which 30,000 and 5,000 reads were generated with the mRNA and BCR libraries respectively.

### scRNA-seq data processing

For all scRNA-Seq experiments, UMI counts were calculated using CellRanger (v.3.1.0) and reads were aligned using mm10 as the reference genome. CellRanger filtered feature sparse matrix counts files from the CD138^+^ splenocyte sample obtained on 35 d.p.i. (HDF5 format) were supplied for unsupervised cell-clustering analysis using the ICGS2 algorithm in the software AltAnalyze using default parameters^3^. ICGS2 delineates non-negative matrix factorization (NMF) clusters and unique marker genes. Cell-type/state labels were initially based on ICGS2 cluster-specific marker gene-set enrichment (GO-Elite) and then further refined with manual curation using literature-associated markers. A small proportion of non-B-lineage cells were identified based on their marker gene profiles and were excluded from the downstream analysis. All scRNA-Seq datasets presented and used for analysis will become publicly available on GEO.

### scRNA-Seq annotation and integration

ICGS2 cell clusters for CD138^+^ splenocytes isolated from 35 d.p.i. were used as the reference for all remaining single-cell experiments presented (supervised classification). Label transfer was performed in the software cellHarmony^4^ using centroid-based classification and a minimum Pearson correlation coefficient of 0.7 on the log2 scaled gene expression counts, to exclude low-quality alignments.

### Calculation of cell cluster frequencies in CD138^+^ splenic subsets

CD138^+^ splenic subsets were flow sorted based on the indicated cell surface markers and processed for scRNA-seq. Cell clusters within the CD138^+^ splenic subsets were identified using the outputs from cellHarmony. The frequency of cell clusters within the mixed group of purified B220^+^CD138^+^CD44^+^CD11a^+^ and B220^int^CD138^+^CD44^+^CD11a^+^ splenocytes were calculated by subtracting the proportion of cell clusters contained within the unmixed B220^−^ CD138^+^CD44^+^CD11a^+^ cells processed using scRNA-seq.

### Differential gene expression analyses

Differential gene expression analysis was performed using cellHarmony between the reference (CD138^+^ splenocytes – 35 d.p.i.) and the query cluster. Genes with an absolute fold > 1.2 and empirical Bayes t-test p-value < 0.05 (Benjamini Hochberg corrected) were considered differentially expressed. A chi-square test was performed to assess differential cell type frequency within.

### Pseudotime trajectory and RNA velocity analyses

Cell-by-gene counts and cell type labels file were filtered for cells from LZ and DZ GC B cell, PC progenitor and distinct PC clusters, and were used as input for Monocle2 pseudotime analysis using default options. LZ GC B cells were indicated as the root of the analysis. The counts data was modeled with negative binomial distribution using Monocle 2 (‘expressionFamily=negbinomial.size’). Monocle 2 was allowed to select its own genes for pseudotime estimation based on differential gene analysis across the filtered B-lineage cells (‘fullModelFormulaStr = ∼Groups’). The reverse graph embedding (RGE) (‘method’ in reduceDimension) method was set to “DDRTree” as recommended^5^. The Monocle2 cell state that contained the maximum number of LZ B cells was set as the “root” state to calculate the pseudotime of all cells.

For RNA velocity analysis, the cell per gene matrix generated from the cell ranger, filtered for LZ and DZ GC B cells, PC progenitors and distinct PC clusters, was provided as an input for Velocyto tool to generate a loom file that contained the spliced and unspliced counts for each gene in each cell^6^. The loom file was then preprocessed using the default parameters described in scVelo package (Version 0.2.5)^7^ and Scanpy (Version 1.9) packages^8^.

### Gene signature score

To calculate signature scores of G2/M genes, an algorithm from our previous study^2^ was adopted to control the variation of quality and complexity of scRNA-seq data of individual cells. To calculate the gene signature score of PCs, this algorithm was modified based on the top 50 upregulated genes analyzed in BMPCs by bulk RNA-seq along with four important PC genes^9^.

### BCR analysis of 5’-end scRNA-seq

Single-cell VDJ sequencing data were processed using Cellranger VDJ (v.3.1.0 – 5 Prime V1; v.6.1.2 – 5 Prime V1.1) reference for mouse from 10x Genomics. Cell barcode associations from CellRanger VDJ and linked gene expression were used for coupling the BCR and transcriptomes, respectively. Cells that did not contain both the BCR and gene expression features were excluded for all downstream clonal analyses. The somatic hypermutation rate and affinity maturation were determined for these cells as described previously^2^. Clones were identified based on cells that have identical V(D)J rearrangements in their heavy and light chain loci (kappa or lambda), including shared amino acid sequences encoded in their *IGHV* CDR3 regions. If a cell contained multiple *kappa* and/or *gamma* chain rearrangements, then the productively rearranged light chain gene was used for clonal identity. This analysis was performed for 22 NP-specific *IGHV* genes and 19 KLH-specific *IGHV* genes (Supplementary table 2). The R code defining the clones is provided at (https://github.com/kairaveet/bcr-clones).

### Statistical analysis

For the biological datasets, we used Prism version 9 (GraphPad) for differential testing. Pairwise statistical tests were performed using a two-tailed Student’s t-test, whereas multicondition comparisons were performed using a one-way ANOVA with Tukey’s multiple comparison test or Kruskal Wallis with Dunn’s multiple comparison test were used based on the distribution, as indicated. For comparison of multiple measures of same samples for more than two or more timepoints, repeated measures two-way ANOVA with Tukey’s multiple comparison test was used.

## Acknowledgments

We thank the expert personnel within Division of Laboratory Animal Resources, the Flow Cytometry Core, the Single Cell Core and UPMC Genome Center at the University of Pittsburgh for their invaluable assistance. We particularly wish to acknowledge Tracy Tabib for help with the scRNA-seq and BCR-seq library preparations and Stuart Hay for setting up the viewer for scRNA-seq datasets. This research was supported by the UPMC ITTC fund and NIH grants U01AI141990, RO1 AI145064.

## Author contributions

H.S. and H.X. conceived and supervised the study. G.K.M.V. and M.Z. performed all experiments. A.R., A.S.C. and A.J. assisted with some of the immunization and adoptive transfer experiments. G.K.M.V. and K.T. performed the computational analysis. L.C.L., P.H., D.C. and K.C. helped with the computational analysis. P.C., J.F. and J.D. helped with some of the data representation. A.J. and L.B. provided critical inputs during the study. N.S. supervised the computational analysis. G.K.M.V., M.Z., H.X. and H.S. wrote the manuscript with input from other authors.

